# High cell-type specificity of eQTLs revealed by single-nucleus analyses of brain and blood

**DOI:** 10.1101/2025.11.26.689911

**Authors:** Martijn Vochteloo, Anoek Kooijmans, Joost Bakker, Roy Oelen, Jelmer Niewold, Dan Kaptijn, Marc Jan Bonder, Monique van der Wijst, Yunfeng Huang, Julien Bryois, Ellen A. Tsai, Lude Franke, Harm-Jan Westra

## Abstract

Identifying causal mechanisms from genome-wide association studies (GWAS) requires an understanding of how disease-associated genetic variants influence gene expression in specific cell types. Here, we present scMetaBrain, a large-scale single-nucleus RNA-sequencing (snRNA-seq) resource derived using 1,260 samples from 785 individuals spanning 10 brain datasets. By analyzing 3.9 million transcriptomes, we identified 19,371 unique expression quantitative trait locus (eQTL) genes (eGenes) at a major cell type level, with the largest number of eQTLs observed in excitatory neurons. Notably, 31% of the eQTLs detected were highly cell-type-specific, with most restricted to excitatory neurons (69%). We compared the eQTLs with bulk RNA-seq datasets across different tissues and with a newly generated single nucleus dataset of 123 donors from peripheral blood mononuclear cells. We observed that differences in eQTL effect sizes between brain cell types are often as large as comparing eQTLs between brain tissue and non-brain tissue from bulk RNA-seq studies. Furthermore, we observe that eQTL effect size agreement was highest for cell types with similar function, even when comparing brain to blood cells. This suggests that that bulk analyses substantially overestimate eQTL agreement, likely due to tissue-level averaging of cellular regulatory effects. Through colocalization, we prioritized 662 genes for 11 brain-related traits and prioritized a single cell type in 68% of genes. Our findings demonstrate that eQTL effects are far more cell-type-specific than previously recognized, underscoring the need to expand single-cell eQTL studies across diverse tissues and cell types to fully capture the regulatory architecture of genetic variants.

## Introduction

Millions of individuals worldwide suffer from brain-related diseases. Understanding these complex genetic diseases requires not only identification of disease-associated genetic variants but also characterization of their functional effects in relevant cell types. Genome-wide association studies (GWAS) have uncovered thousands of loci associated with neurological and psychiatric disorders^1^, yet most associated variants reside in non-coding regions, making it difficult to pinpoint their functional consequences. To bridge this gap, expression quantitative trait locus (eQTL) studies provide a powerful approach for linking genetic variation to changes in gene expression^2–8^, thereby revealing regulatory mechanisms underlying disease.

Multi-tissue eQTL resources such as GTEx^9–12^ have proven invaluable for the interpretation of GWAS loci, and these have been followed up by more specialized single-tissue resources such as those created by the eQTLgen consortium^4^ in blood and PsychENCODE^5^, AMP-AD^13^, and MetaBrain^14^ in brain. These studies have shown that while tissue specific eQTLs exist, there is high sharing of eQTLs across different tissues, including different brain regions^15^. This may complicate the interpretation of disease-associated loci. We hypothesized that this apparent sharing might be because most large eQTL studies mainly rely on bulk transcriptomics, which lacks cell type resolution, potentially obscuring cell-type-specific regulatory effects. Recent efforts to map eQTLs using single-nucleus transcriptomics derived from brain tissues have revealed additional eQTLs that were not observed in whole tissue^7,8,16,17^, despite their lower sample size. Apart from driving hypotheses about the mechanisms behind neurological disease, these resources also pinpoint putative driver cell types. However, these studies do not yet have a sufficient sample size to maximize the number of significantly detected eQTLs. This is especially true for highly disease-relevant but low abundance cell types in brain such as microglia. Moreover, increased sample sizes can improve eQTL effect size estimation, which consequently improves the performance of downstream analyses such as colocalization.

In this study, we present a well powered, cell-type-specific eQTL resource for studying the gene regulatory effects of genetic variants associated with neurological diseases. We combined and rigorously harmonized single-nucleus transcriptomic and genomic data from 10 brain datasets, encompassing 1,260 brain snRNA-seq samples from 785 individuals, mostly from cortical brain areas (**Figure 1A**). We mapped eQTLs in eight major brain cell types (**Figure 1B**) and compared our results to those from large bulk-tissue datasets across multiple tissues and a newly generated single-nucleus dataset from blood. We show that the functional context of the cell defines genetic regulation of gene expression rather than the local tissue-environment (**Figure 1C**). Furthermore, we conducted systematic colocalization analyses to detect shared genetic effects between eQTLs and 11 brain-related GWAS traits (**Figure 1D**), which prioritized 662 disease-relevant genes and their potential cell type context. These findings highlight the importance of expanding single-cell eQTL studies across diverse tissues to fully capture the regulatory architecture of genetic variants. This will ultimately advance our understanding of neurological disease mechanisms and improve therapeutic target discovery.

**Figure 1.**
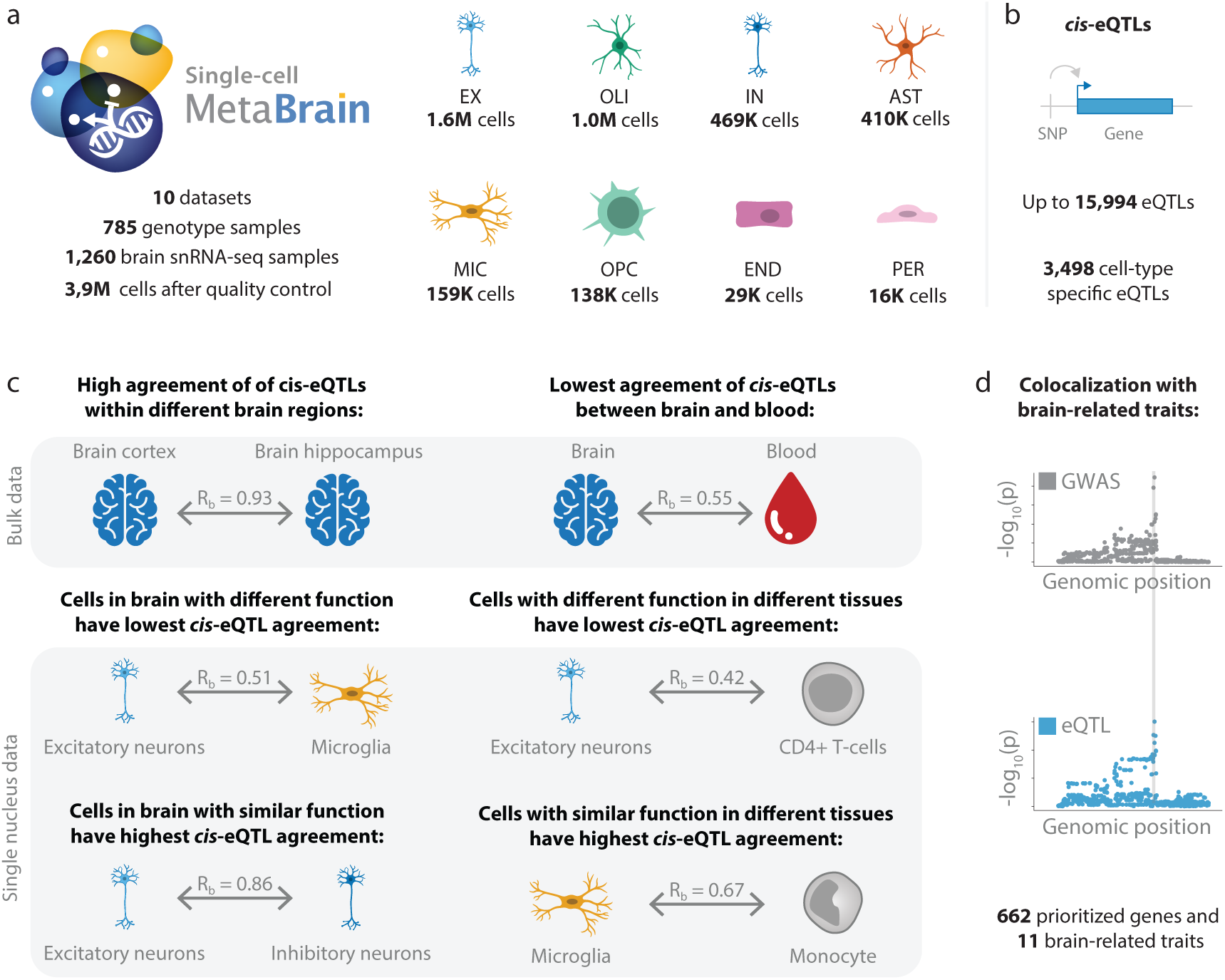
Overview of the study. **a,** We processed publicly available RNA-seq and genotype data from 10 different datasets consisting of 1,260 snRNA-seq amples and 785 genotype samples. We created cell-type-specific eQTL datasets for each of the eight major brain cell types, and **b,** performed *cis*-eQTL analyses. **c,** By comparing to eQTLs from bulk and single cell datasets in brain and blood, we determined the cell-type-specificity of *cis*-eQTLs. **c,** Finally, we prioritized genes for brain-related phenotypes using colocalization. Cell type images are created with BioRender.com.

## Results

### Harmonizing single-nucleus datasets for eQTL analysis

We integrated 10 single-nucleus eQTL datasets derived predominantly from the human prefrontal cortex area of the brain into the ‘scMetaBrain’ resource to maximize the statistical power to detect eQTLs (**Figure 1** and **Supplementary Table 1 and Supplementary Table 2**). In total, we processed 1,370 snRNA-seq and 1,210 genotype samples from the AMP-AD consortium (Mathys^18^, Sun^16^, Zhou^19^, Cain^20^, Fujita^8^, and Huang^21^), the PsychENCODE consortium (Ruzicka^22^), and Roche (RocheAD^7^, RocheMS^7^, and RocheColumbia^7^). For our analysis, we focused on eight major cell types: excitatory neurons (EX), inhibitory neurons (IN), microglia (MIC), astrocytes (AST), oligodendrocytes (OLI), oligodendrocyte precursor cells (OPC), endothelial cells (END), and pericytes (PER). After quality control (QC), normalization, sample integration selecting samples from duplicate snRNA-seq and genotype samples, genotype imputation, and doublet removal (**Methods**), 1,260 snRNA-seq samples for up to 785 unique individuals remained, with final sample sizes depending on the cell type (**Supplementary Table 2**). We obtained gene expression data for between 20,922 (OPC) and 27,766 genes (EX), 61−71% of which were protein-coding. Total cell counts ranged between 16,349 (PER) and 1,609,788 (EX) cells (**Supplementary Table 2**). We tested genotypes for 7,242,888 variants (minor allele frequency (MAF)>5%, imputation quality score>0.3) of which 80% were single nucleotide polymorphisms (SNPs) and the remainder indels.

### Indels are more likely to be eQTLs but are located further from the transcription start site (TSS)

In each cell type, we conducted a *cis*-eQTL analysis on common variants (MAF>5%) located within 1 million-base pairs (Mb) of the TSS of each expressed gene using linear regression. Combined over all cell types, we observed a significant eQTL (q-value<0.05) in one or more cell types for 19,371 genes (out of 29,030 total tested genes), with numbers of significant eQTL genes (eGenes) ranging from 136 (PER) to 15,994 (EX) (**Figure 2A** and **Supplementary Table 3**). We observed a high correlation between the number of eQTLs detected and the total number of nuclei per cell type (Pearson r=0.93; **Figure 2A**). This indicates that having larger numbers of nuclei enhances gene expression quantification and eQTL discovery by enabling inclusion of genes with lower expression and increasing power to detect eQTLs with smaller effect sizes.

**Figure 2.**
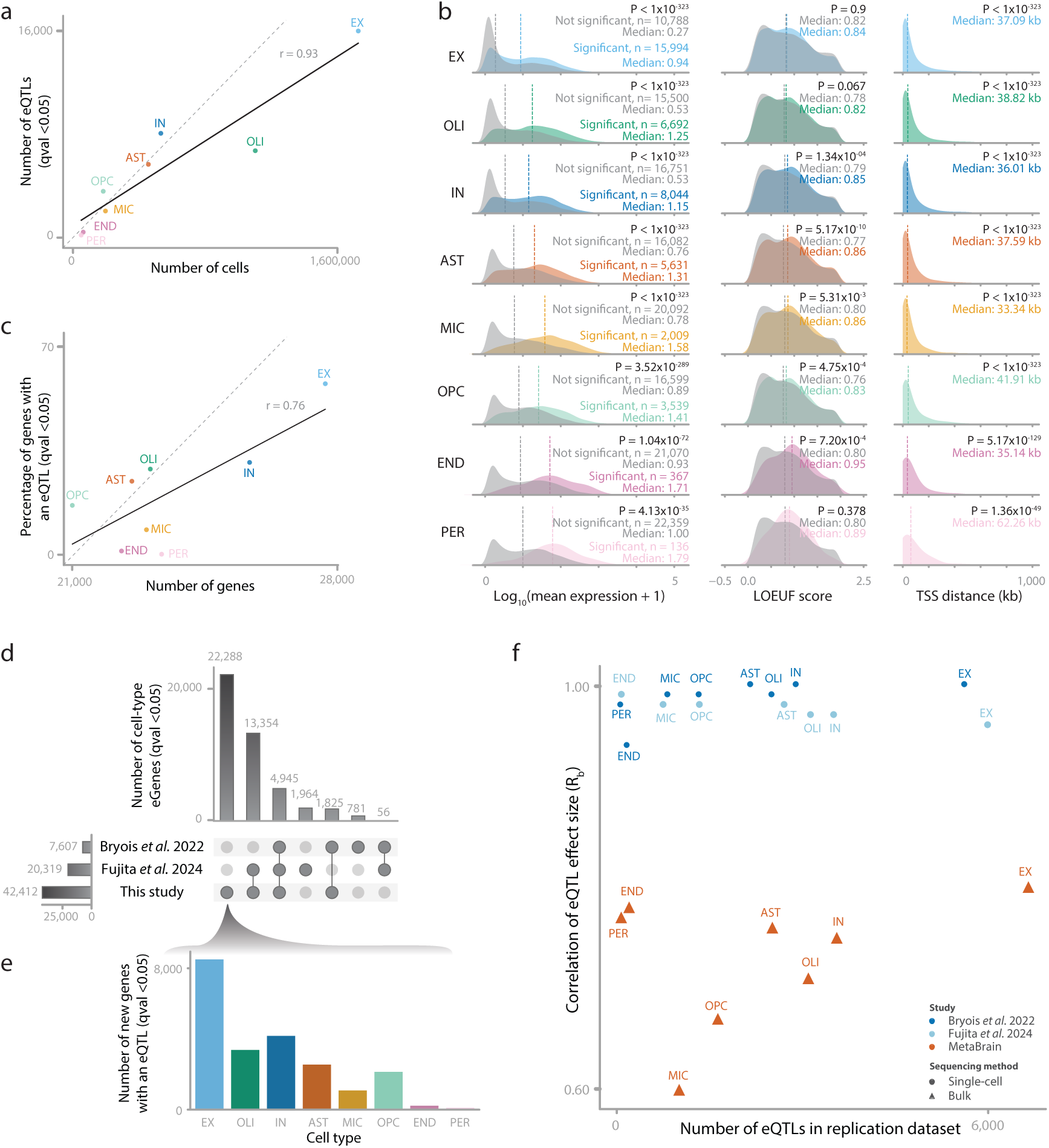
*cis*-eQTLs and their characteristics. **a**, Number of cells (x-axis) versus the number of *cis*-eQTLs (y-axis) per cell type. **b**, Comparison of characteristics between eQTL genes and non-eQTL genes. P-values were calculated using a two-sided t-test between significant and non-significant genes. Shown are the differences in mean gene expression levels (left), LOEUF scores (middle), and the distance between the most significant variant and the TSS per gene (right). Vertical dotted lines indicate the median of the distribution for the eGenes (colored) versus the non-eGenes (grey). **c**, Number of expressed genes (x-axis) versus the proportion of genes with an eQTL (y-axis) per cell type. **d, e,** Number of overlapping cell type-eGene combinations between this study and Bryois *et al.* and Fujita *et al.* (**d**) and the number of newly detected eQTLs per cell type (**e**). **f**, Correlation of the effect sizes of *cis*-eQTLs per cell type (discovery) in the corresponding cell type in the brain-derived single-nucleus eQTL studies from Bryois *et al.* (dark blue, replication) and Fujita *et al.* (light blue, replication), as well as the bulk RNA-seq eQTLs from MetaBrain Cortex-EUR in which each discovery cell type is compared to the same bulk eQTL signal (orange, replication). Each dot is an eQTL replication dataset. X-axis indicates the number of eQTLs that are significant in the discovery dataset and overlap with the replication dataset.

We next investigated the characteristics of the eQTL genes, their lead variants, and lead variant−gene combinations. Investigating gene characteristics, we observed that eGenes showed significantly higher expression than non-eGenes within each cell type (two-tailed t-test p-value<1.08×10^-35^; **Figure 2B**). In all cell types except EX, OLI and PER, eGenes showed lower evolutionary constraint as indicated by higher LOEUF-scores^23^ compared to non-eGenes (two-tailed t-test p-value<5.31×10^-3^; **Figure 2B**), similar to earlier observations^7^. We reasoned that the lack of significance in EX, OLI and PER was most likely due to the relatively high (EX, OLI) or low (PER) number of significant eGenes compared to the number of tested non-eGenes.

Next, we investigated the characteristics of the lead variants for each eGene, limiting the analysis to eGenes (q-value<0.05) within each cell type. For 29.9% of *cis*-eQTLs, the lead variant was an indel (12,687/42,412), while only 20% of all variants tested were indels (1,438,427/7,242,888), indicating that indels are significantly enriched for regulating gene expression (two-sided Fisher exact test p-value<3.5×10^-323^; **Supplementary Table 3**). This observation could not be explained by differences in statistical power between indels and SNPs, because we did not observe a significant difference in eQTL effect sizes between both types of variants in any of the tested cell types (two-tailed t-test p-value>0.08). The average indel length was 3.3 bases and was mostly consistent between cell types (two-tailed t-test p-value>0.11), except between IN and OLI (two-tailed t-test p-value=0.01). Only in OLI did we observe that eGenes with an indel as lead variant had significantly lower average expression than eGenes with a SNP as lead variant (two-tailed t-test p-value=0.04). We also observed that eGenes with an indel as lead variant in EX, IN, and OLI had significantly lower LOEUFs compared to eGenes with a SNP as lead variant (two-tailed t-test p-value<1.2×10^-3^). Finally, we observed that the lead eQTL variants were relatively proximal to the TSS (median distance: 33−42 kilobases (kb); **Figure 2B**) in all cell types except PER, where the variant was considerably further from the TSS (62 kb). This may suggest lower confidence for eQTLs in PER, most likely due to limited numbers of these cells. We further observed a significantly higher absolute TSS distance for eQTLs with an indel as lead variant as compared to eQTLs with a SNP as lead variant for all cell types (two-tailed t-test p-value<0.02), except for PER cells (two-tailed t-test p-value=0.10).

Overall, the characteristics of the *cis*-eQTLs we observed were consistent with our previous findings in blood^4^ and brain^14^, suggesting that our approach identifies valid eQTLs. However, although indels appeared to be enriched for eQTLs, we noticed that these eQTLs had a greater TSS distance and generally lower gene expression, warranting follow-up research on these systematic differences.

### scMetaBrain shows high agreement with previous single-cell studies while also identifying additional eQTLs

To further test the validity of the identified eQTLs, we determined replication with previous single-nucleus studies. The scMetaBrain resource includes datasets that have previously been used for *cis*-eQTL discovery in two separate studies: Bryois *et al.*^7^ (n=192) and Fujita *et al.*^8^ (n=424). Combined over all cell types, we report 42,412 significant eQTLs (i.e. unique cell type-eGene pairs, q-value<0.05), which is a 5.6-fold increase compared to Bryois *et al.* (34,805 additional pairs) and a 2.1-fold increase compared to Fujita *et al.* (22,093 additional pairs; **Figure 2D**). A total of 22,288 cell type-eGene pairs (52.6%) were not previously reported in either study, most in EX (8,517; **Figure 2E**). For each of the eight cell types, we compared the direction and effect size of our eQTLs with the findings reported by these two earlier studies (**Figure 2F** and **Supplementary Table 4**). Combined over all cell types, we were able to overlap 15,390 (36%) of our 42,412 significant eQTLs with Bryois *et al.* and 17,349 (41%) with Fujita *et al.* In most cases where we were unable to overlap the eQTL (72.5% in Bryois *et al.* and 74.9% in Fujita *et al.*), the gene was not present in the reported summary statistics, potentially due to the lower number of cells in the respective studies. For the overlapping effects, we determined their rate of agreement using three different measures: allelic concordance (AC), correlation of effect sizes (*R_b_*), and the proportion of true positives (*π_1_*) between each of the corresponding cell types. We observed high agreement with both the Bryois *et al.* (AC>95%, *R_b_*>0.94, *π_1_*>0.73; **Figure 2F**) and Fujita *et al*. studies (AC>99%, *R_b_*>0.96, *π_1_*>0.83; **Figure 2F**). We note that the *π_1_* estimates were somewhat lower than AC and *R_b_*, perhaps because our analysis had a larger sample size. As such, when replicating effects in a smaller cohort, we considered *π_1_* estimates as the lower bound estimate of sharing: the high AC and *R_b_* estimates indicate that the additional eQTLs in our analysis have similar directions and effect sizes even when they were not significant in the previous studies, which suggests that these studies were likely underpowered to detect these effects with confidence.

### Rate of agreement with whole tissue eQTLs depends on cell type composition of those tissues

Having concluded that the eQTLs in our study showed good replication with those from previous single-nucleus brain studies, we aimed to investigate their tissue- and cell-type-specificity. To do this, we first compared the eQTLs for each of the cell types with eQTL summary statistics from our previous bulk RNA-seq MetaBrain study^14^ and with bulk eQTL summary statistics from other tissues from the GTEx consortium^12^. We then zoomed in on cell type differences by first comparing the eQTLs between the different brain cell types in our study and then by investigating our eQTLs in single blood cell types.

To investigate the tissue-specificity of the observed brain cell type eQTLs, we first evaluated replication of the scMetaBrain eQTLs using eQTLs from our previous whole-tissue brain cortex study (MetaBrain Cortex-EUR^14^, n=2,683; **Supplementary Table 4**). Since MetaBrain only reported eQTLs for SNPs and protein-coding genes, we limited the replication to the 18,581 overlapping eQTLs (representing 64.7% of scMetaBrain eQTLs). Combined over all cell types, the AC was 84%, indicating that in 16% of the cases the direction of effect in the single-nucleus data was opposite to that in the bulk data. Per cell type, we observed moderate rates of agreement (AC>78%, *R_b_*>0.60, *π_1_*>0.54; **Figure 2F** and **Supplementary Table 4**). The *R_b_* and AC measures were lowest in MIC (*R_b_*=0.59, AC=0.78) and OPC (*R_b_*=0.67, AC=0.80). Excluding the END and PER cell types, which had few overlapping eQTLs (n<201), we observed a correlation on the number of included cells for the *R_b_* (Pearson r=0.69) and AC measures (Pearson r=0.67), which could indicate these measures are underestimated for MIC and OPC. On the other hand, compared to the neuronal cell types, MIC and OPC have low abundances in the brain and may therefore also contribute less to the overall whole-tissue transcriptomics signal, which also might explain the lower agreement estimates.

To further test this, we next compared our results with eQTLs from the other brain regions from the MetaBrain study (**Supplementary Figure 1** and **Supplementary Table 4**), we found that the eQTL agreement of brain cell types differed between brain regions, with specific cell types showing the highest agreement in specific brain regions. For instance, EX and IN showed highest agreement in cortex (*R_b_* =0.80 and *R_b_*=0.75, respectively), whereas OLI and MIC showed highest agreement in spinal cord (*R_b_*=0.79 and *R_b_*=0.72, respectively). We extended this comparison to other tissues by comparing the scMetaBrain eQTLs to bulk eQTLs in 49 distinct tissues from GTEx^2^ (**Supplementary Figure 1** and **Supplementary Table 4**). Here we again observed large differences in agreement between tissues, but highest rates of agreement in those tissues that are mainly composed of those cell types: cortex showed the highest agreement for EX (*R_b_*=0.70), followed by other brain regions such as cerebellum and cerebral tissues. Additionally, spinal cord (cervical c-1) showed the highest agreement with OLI (*R_b_* =0.74) and cerebellum showed the highest agreement with IN (*R_b_*=0.68).

Finally, MIC showed the highest agreement with whole blood (*R_b_*=0.61), while all other cell types had considerably lower similarity to whole blood (average *R_b_*=0.48). These similar observations between GTEx and bulk MetaBrain make it less likely that the observed *R_b_* estimates are highly dependent on the number of included cells or tested genes per scMetaBrain cell type. Overall, we observed that the *R_b_* values for the comparisons with bulk tissue eQTLs were considerably lower than those observed when comparing bulk brain tissue eQTLs. For instance, whereas the best powered scMetaBrain eQTL cell type, EX, had *R_b_*estimates ranging between 0.36 and 0.7, the bulk comparisons between MetaBrain and GTEx^14^ ranged between 0.68 and 0.96, suggesting higher specificity of single-nucleus eQTLs compared to bulk eQTLs.

### eQTLs are more often shared between biologically similar cell types, even across tissues

To investigate the differences between cell types, we first determined to what extent eQTLs were specific to one cell type in the brain. We observed that 8,323 (43%) of the 19,371 eGenes were significant in just one cell type. This number decreased to 3,498 (31%) when limiting the comparison to 14,044 genes that were expressed in each of the cell types (<90% zero values in all cell types), 11,225 of which were eQTL genes (**Supplementary Table 3**). The EX cell type had the highest number of eQTL genes that were not observed in another cell type (2,420/3,498=69%) and PER the lowest (0), with a clear correlation between the number of cells per cell type and the number of unique eQTLs (Pearson r=0.9). To reduce this cell-count effect, we therefore next quantified the number of unique genes after grouping the eQTLs from the neural lineage (EX and IN), the glial lineage (AST, OPC, and OLI), and endothelial-like cells (END and PER), keeping MIC as a separate group (**Supplementary Table 3**). This showed that 44% of eQTL genes (4,911/11,225) that are observed in one cell type lineage are not observed in another.

We next determined the agreement of eQTLs between the cell types within our study (**Figure 3A and B**, **Supplementary Figure 1**, **Supplementary Table 4**). We observed moderate agreement when averaged over all cell types (AC=75%, *R_b_*=0.65, *π_1_*=0.54, percentage of replicating effects with Benjamini-Hochberg false discovery rate (BH-FDR)<0.05=41.7%; **Supplementary Table 4**). The *π_1_* estimates and average proportion of significantly replicating *cis*-eQTLs (BH-FDR<0.05 over overlapping eQTLs) were affected by the number of cells measured (Pearson R=0.64 and 0.63, respectively), with cell types with more cells being more distinct. Notably, the R_b_ averaged over all cell types (R_b_=0.65; **Supplementary Table 4**) was comparable to the average R_b_ obtained when comparing bulk MetaBrain cortex brain region eQTLs with non-brain tissue eQTLs in GTEx (average R_b_=0.67; excluding cell-lines, reproductive tissues and whole blood; **Figure 3A**; results from de Klein *et al.*, 2023). This suggests that differences between cell types in the brain are on average as large as comparing brain tissue with non-brain tissues. The average rates of agreement over all cell types (AC=75%, *R_b_*=0.65, *π_1_*=0.54) were also lower than the averages calculated over the comparisons with the bulk MetaBrain cortex eQTLs (AC=84%, *R_b_*=0.73, *π_1_*=0.79; **Figure 3B**, **Supplementary Table 4**). This suggests that single-nucleus eQTLs are more specific to their respective cell types than to their originating tissue. Compared with earlier work by Fujita *et al.*^8^, we found more eQTLs to be cell-type-specific (lowest AC=0.6 compared to 0.88), indicating that increased sample size yields additional cell-type-specific effects. Finally, we found that biologically similar cell types showed higher agreement than biologically different cell types. For instance, two pairs of cell types that showed notably higher agreement were EX and IN (AC=90%, *R_b_*=0.88, *π_1_*=0.79) and END and PER (AC=91%, *R_b_*=0.82, *π_1_*=0.48), with other pairs of cell types showing notably lower agreement (AC<84%, *R_b_*<0.75; **Figure 3B**). In terms of *R_b_*, the most distinct pair of cell types was END and EX (AC=65%, *R_b_*=0.51, *π_1_*=0.42), while the most distinct individual cell type was MIC (average over all comparisons: AC=73%, *R_b_*=0.58, *π_1_*=0.54; **Figure 3B**). The high specificity of microglia eQTLs compared to other brain cell types, coupled with their low frequency in whole tissue, suggests that many microglia-specific eQTLs may have been missed in bulk eQTL studies and remain to be discovered. Since microglia are known to play an important role in diseases such as Alzheimer’s disease (AD)^24^ and multiple sclerosis (MS)^25^, we expect that especially microglia-specific eQTLs derived from single-nucleus data may help explain additional GWAS loci.

**Figure 3.**
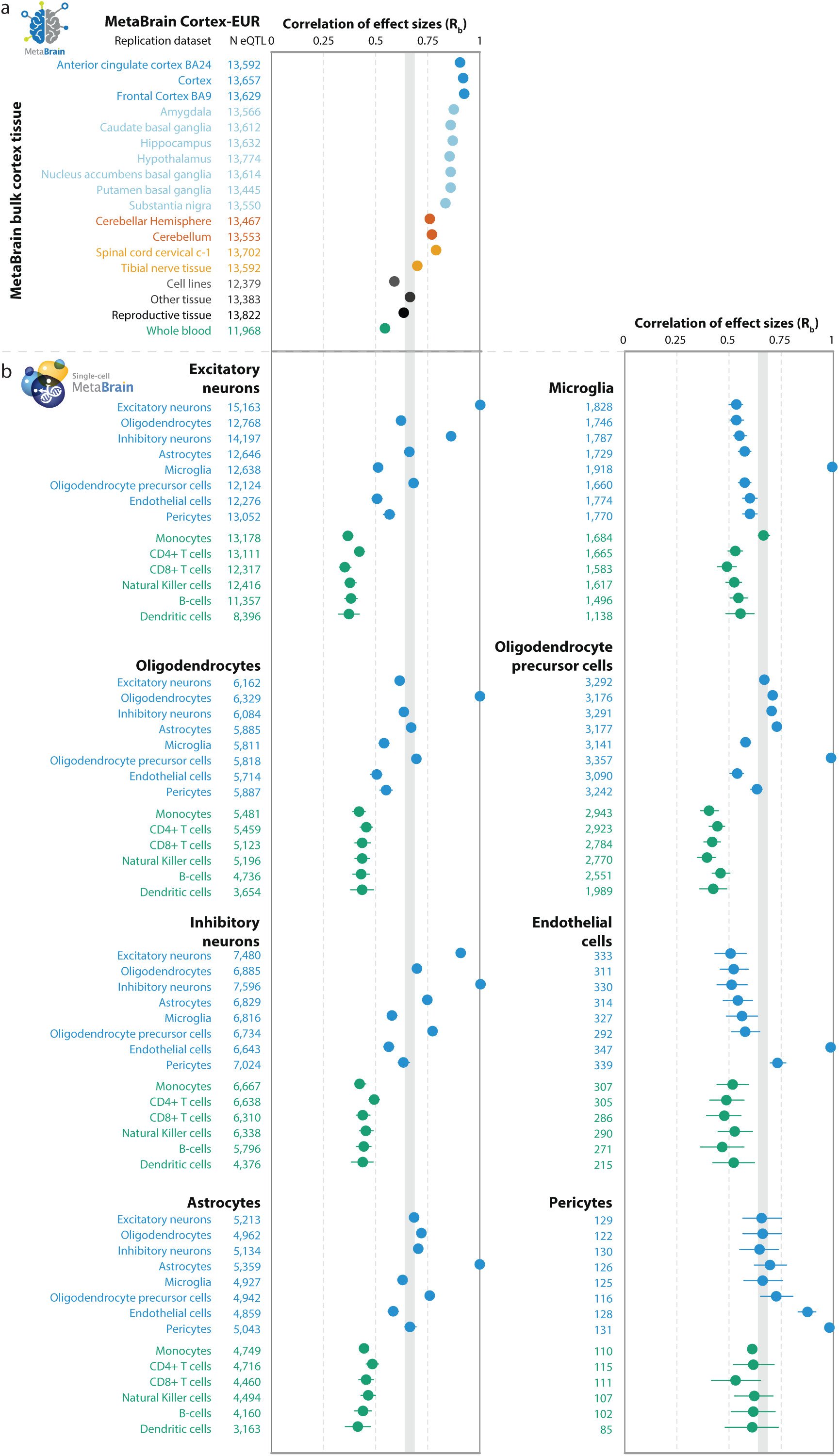
Tissue- and cell-type-specificity of eQTLs. Correlation of effect sizes (R_b_) between *cis*-eQTLs observed in the discovery eQTL dataset and different replication eQTL datasets. Each dot is the R_b_ of a separate eQTL agreement analysis. Horizontal lines indicate the 95% confidence interval. Each block of dots indicates a single replication analysis, where the discovery cell type is indicated in bold text. **a,** For reference, we include a comparison of the MetaBrain bulk Cortex-EUR brain region eQTL dataset with GTEx tissues, derived from de Klein *et al.* 2023. The grey vertical bar indicates the average agreement of MetaBrain bulk cortex eQTLs in an average of non-brain bulk-tissue eQTLs in GTEx. **b,** For scMetaBrain, we compared brain cell type eQTLs between scMetaBrain (cell types indicated in blue) and with cell types from PBMC single-nucleus data (cell types indicated in green). eQTLs observed in single cell samples from the same tissue show a similar rate of agreement to that found when comparing bulk brain to a random non-brain bulk tissue. Comparisons with other eQTL datasets from brain and non-brain tissues can be found in **Supplementary Figure 1**.

Our comparisons among GTEx tissue eQTLs showed that brain cell type eQTLs predominantly replicate well in brain tissues, except for microglia, which replicated best in whole blood. Therefore, to finalize our investigation of the cell-type-specificity of our eQTLs, we compared our eQTLs to those from a single-nucleus blood dataset that we recently generated (manuscript in preparation, n=123, **Supplementary Table 4**). In short, we mapped single-nucleus eQTLs in six blood types: CD4+ T-cells, monocytes, natural killer cells, CD8+ T-cells, B lymphocytes, and dendritic cells. Over all comparisons, we were able to overlap the same variant-gene combinations for 74.6% of all brain eQTLs (q-value<0.05). Brain cell type eQTLs were distinct from blood cell type eQTLs, with overall low averages of agreement (AC=63.54%, *R_b_*=0.48, *π_1_*=0.27). AC and *R_b_*estimates were similar to our comparison of scMetaBrain and whole blood from GTEx (average AC=68.93%, average *R_b_*=0.49), while due to sample size differences, the average *π_1_* was higher in the GTEx comparison (*π_1_*=0.62; **Supplementary Table 4**). However, when comparing individual cell types, the immune-related cell type MIC (*R_b_* =0.56) and vascular cell types such as PER (*R_b_* =0.62) and END (*R_b_* =0.52) showed relatively higher average agreement with all blood cell types compared to other brain cell types (*R_b_*<0.45; **Figure 3B**). We observed the strongest agreement between brain and blood cell types for MIC and monocytes (AC=79%, *R_b_*=0.67, *π_1_*=0.56; **Figure 3B**), highlighting the similarity in the genetic regulation of gene expression between these immune-related cell types even across tissues. This relatively strong overlap suggests that eQTLs are more likely to be shared between cell types with similar biological functions, rather than being strictly tissue specific.

### Shared genetic effects between cis-eQTLs and brain-related traits and diseases

Having observed high cell type specificity of eQTLs between brain cell types, we next investigated whether this also affects colocalization of GWAS associations for brain-related phenotypes and diseases. We tested four neurodegenerative disorders (Parkinson’s disease (PD)^26^, AD^24^, MS^25^, and amyotrophic lateral sclerosis (ALS)^27^), four psychiatric disorders (schizophrenia (SCZ)^28^, major depressive disorder (MDD)^29^, attention deficit hyperactivity disorder (ADHD)^30^ and bipolar disorder (BPD)^31^), and three other brain-related traits (epilepsy^32^, intelligence^33^, and years of schooling (YOS)^34^). Combined over all traits, we tested colocalization between eQTL and GWAS signals for 6,371 eGenes and found colocalization for 1,217 cell type-eQTL-trait combinations (posterior probability for colocalizing signals (PPH4)≥0.8) for 662 distinct genes (10.4%; **Figure 4A** and **Supplementary Tables 5 and 6**). We found colocalization with 3 genes for ADHD, 60 for AD, 6 for ALS, 27 for BPD, 1 for epilepsy, 131 for intelligence, 71 for MDD, 38 for MS, 95 for PD, 106 for SCZ, and 230 for YOS (**Figure 4A** and **Supplementary Tables 5 and 6**). Over all diseases, there was a high correlation between the number of tested genes and the number of colocalizing genes (Pearson r=0.987; **Figure 4B**, **Supplementary Table 6**), suggesting either the statistical power of the GWAS study or the polygenicity of the phenotype determines the total number of colocalizing genes.

**Figure 4.**
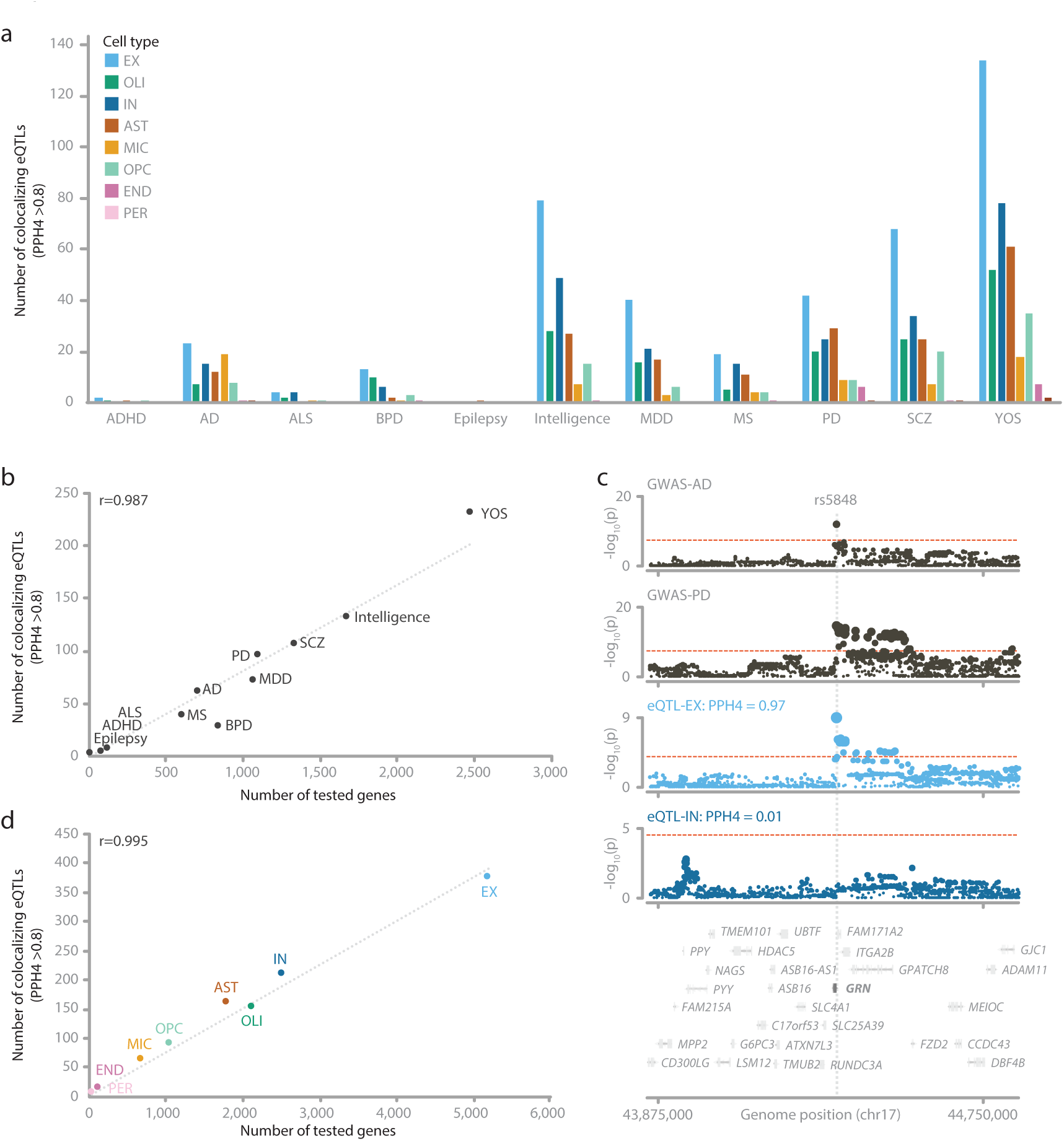
Colocalization analysis of brain-related traits. **a**, Number of colocalizing (PPH4 > 0.8) eQTLs for 11 brain-related traits, per cell type. **b,** correlation between number of tested eQTL genes for each phenotype and the number of colocalizing genes. **c,** Region plots showing an example of a shared, colocalizing signal for AD and PD in the *GRN* locus, that is only observed in the EX cell type. Red dotted line indicates significance threshold for GWAS (p<5×10^-8^) or eQTL (q-value<0.05). **d,** correlation between number of genes tested in each cell type and the number of colocalizing genes.

Averaged over the phenotypes, 29% of the tested GWAS loci (as defined by LocusBreaker^35^) contained at least one colocalizing eQTL, with AD having the highest proportion of colocalizing loci (60%). There was little sharing between the tested phenotypes with 89 genes colocalizing with multiple GWAS phenotypes. Three pairs of phenotypes had more than 10 shared genes: YOS and intelligence (n=34), YOS and PD (n=28), and YOS and SCZ (n=14; **Supplementary Table 6**). Between neurodegenerative diseases, AD and PD shared 8 colocalizing genes. An example of this is for instance the T allele of SNP rs5848 (chr17:44,352,876:C,T), which was previously associated with increased risk for both AD^24^ (p=1.7×10^-12^) and PD^26^ (p=2.18×10^-15^). This allele significantly decreases the expression of nearby gene *GRN* specifically in EX cells (Z-score=-6.08; p=1.2×10^9^) and shows only limited eQTL signal the other cell types (e.g. OLI p>1×10^-4^; MIC p>2×10^-3^; **Figure 4C and Supplementary Figure 2; Supplementary Tables 5 and 6**). *GRN* encodes the anti-inflammatory progranulin, which is subsequently cleaved into granulin, which, in contradiction to its precursor, is involved in inflammatory processes in the brain^36^. Repression of *GRN* could therefore lead to increased inflammatory processes observed in both neurodegenerative diseases. Decreased expression for *GRN* was described previously for AD and PD^37^, although this analysis did not focus on specific cell types. Our results suggest that the AD and PD loci decrease expression of *GRN* specifically in neuronal cells. While for AD we did not identify additional colocalizing genes in the *GRN* locus, we identified 18 additional genes for PD (**Supplementary Table 6**): *CRHR1, FAM215B, ITGA2B*, *KANSL1*, *KANSL1-AS1*, *LRRC37A, LRRC37A2, MAPT*, *MAPT-AS1, MAPT-IT1*, and *PLEKHM1* and seven non-coding genes. Different genes in this locus showed colocalization in different (combinations of) cell types. For instance, the *ITGA2B* eQTL was significant in both neuronal subtypes but only showed colocalization in IN (PPH4=0.97), although it had a relatively high eQTL p-value (p=1.6×10^-6^), whereas the more significant eQTL effect in EX cells (p=4.27×10^-12^) did not show colocalization (PPH4=0.01). Similarly, *PLEKHM1* showed colocalization in IN cells (PPH4=0.94), and OLI (PPH4=0.98) but not in EX cells (PPH4=0.001) and was not a significant eQTL in the other cell types. This could suggest pleiotropy within the *GRN* locus in PD, with the risk alleles for PD affecting different genes in different cell types.

We next determined the cell-type-specificity of the colocalizations, which at first glance, were highly cell-type-specific: overall 68.2% of the colocalizing signals were specific to a single cell type **(Supplementary Table 6)**. Comparing cell types over all diseases, the number of total colocalizing genes and the number of genes colocalizing in a single cell type were highly correlated (r=0.995; **Figure 4D**) with the number of tested genes within each cell type. These proportions and correlations did not markedly change when focusing on the 14,044 genes expressed in all cell types and by grouping the cell types by lineage (**Supplementary Table 6**), making this observation less likely due to differences in statistical power. These results suggest that the likelihood of finding a colocalizing gene in a cell type highly depends on the power of both the eQTL and GWAS studies, but that individual colocalizing eQTLs are likely highly cell-type-specific.

We especially observed high cell-type-specificity for AD. Out of 60 colocalizing genes, eQTLs in EX cells showed the highest number of colocalizations (n=23, 38%), which was expected due to the large number of eQTLs and cells for this cell type. However, the cell type with the second highest number of colocalizations was MIC (n=19, 32%; 14 LocusBreaker loci), even though the number of cells and consequently tested eQTL genes in this cell type was much lower than EX, IN and OLI. This also became apparent when correlating the number of tested genes with the number of colocalized genes where MIC was a clear outlier (**Figure 5A**; **Supplementary Figure 3** and **Supplementary Table 6**). Furthermore, 14 of the 19 genes colocalizing in MIC (from 11 LocusBreaker loci) did not colocalize in any other cell type (*AC090559.1, BIN1, CASS4, CCDC6, CYP27C1, EPHA1-AS1, FERMT2, INPP5D, NDUFAF6, PICALM, RABEP1, RASGEF1C, USP6NL, and ZYX)*, even though 13 of those genes were also a significant eQTL in another cell type (**Figure 5B**). This list of genes includes several notable genes for AD pathophysiology. For instance, after the *APOE* locus, the locus containing *BIN1* and *CYP27C1* is known to have the second strongest association with AD^24^. *BIN1* and *CYP27C1* had significant eQTLs across multiple cell types, including EX (*BIN1* p=6.5×10^-14^; *CYP27C1* p=7.2×10^-6^), IN (BIN1 p=2.9×10^-6^; *CYP27C1* p=2.2×10^-5^), and OLI (BIN p=9.7×10-6; *CYP27C1* not significant), but did not show colocalization in those cell types (PPH4<0.03; **Figure 5C**). In MIC, however, the AD risk allele SNP rs6733839 (T-allele; p=6.48×10^-90^; chr2:127,048,027:C,T) increased the expression of both *BIN1* (p=8.98×10^-16^; z-score=8.04; PPH4=0.99) and *CYP27C1* (p=1.71×10^-8^; z-score=5.64; PPH4=0.99). *BIN1* regulates inflammatory processes in the brain^38^ and different isoforms of *BIN1* have been found to be associated with tau tangle accumulation^39^. *CYP27C1* encodes a Cytochrome P450 family enzyme and is involved in lipid metabolism, but its involvement in AD is less well described. Another example of a MIC specific colocalization with AD is for the *FERMT2* gene. While showing strong colocalization (PPH4=0.96) in MIC, this gene showed distinct significant association signals but did not show colocalization (PPH4<0.01) in the other cell types with a significant eQTL, including EX (p=2.03×10^-8^) and OLI (p=1.5×10^-12^; **Figure 5D**). In MIC, the AD risk allele A for SNP rs17125924 (chr17:52,857,268:A,G) decreased the expression of *FERMT2* (p=2.45×10-7; z-score=-5.16). *FERMT2* was recently found in a siRNA screen as a modulator of amyloid precursor protein metabolism, with inhibition of the gene showing an increase in cerebrospinal fluid Aβ peptide levels of AD cases^40^. Together these results show that many AD association signals colocalize with eQTLs in brain cell types, with a disproportionate number being specific to MIC cells. This further cements the importance of this cell type for AD pathophysiology.

**Figure 5.**
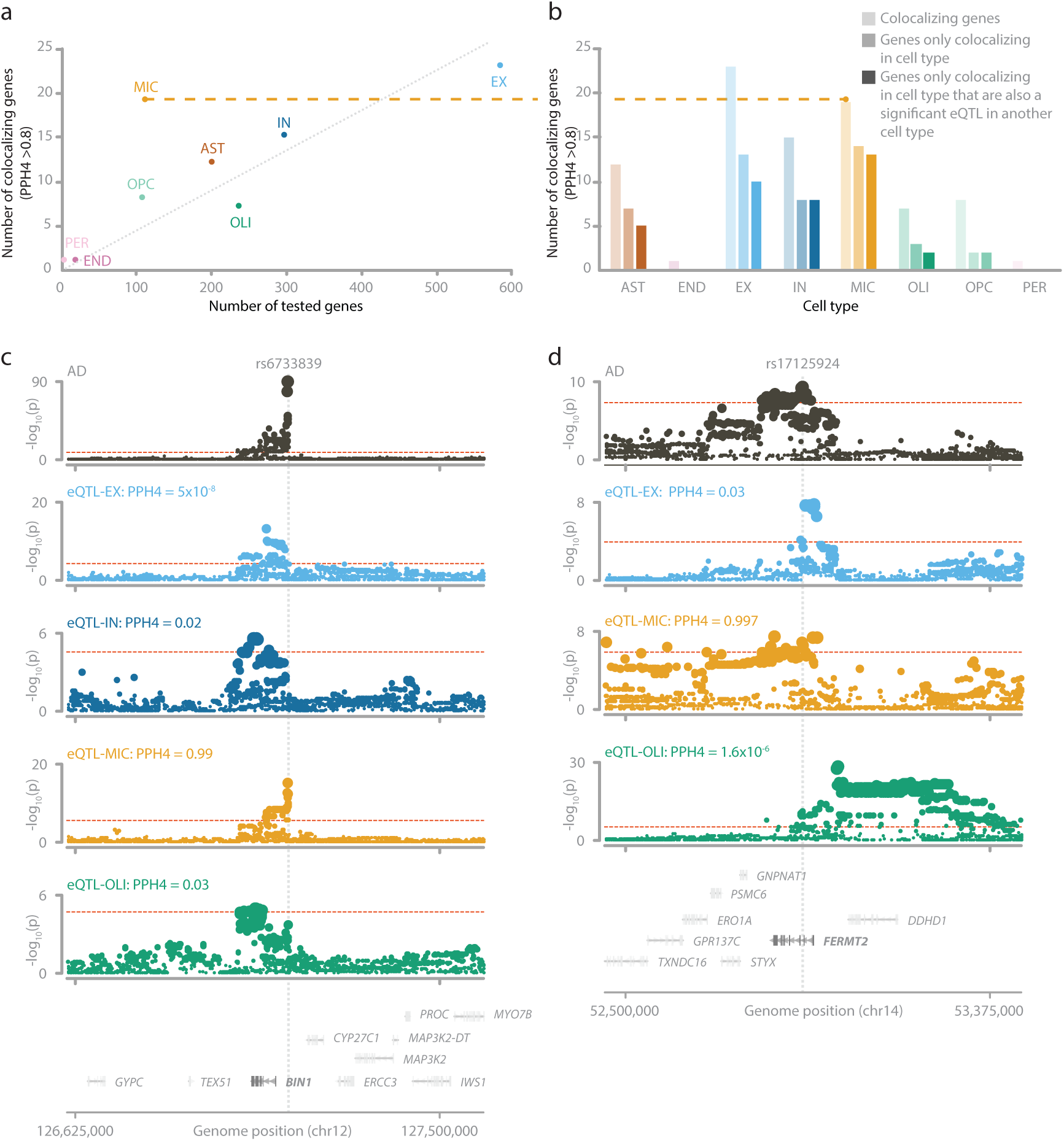
Colocalization results for AD. **a**, Comparing tested genes and colocalizing genes per cell type, the MIC cell type is an outlier, showing a larger number of colocalizing genes than expected from the number of tested eQTL genes. **b,** Many of the colocalizing genes per cell type are unique to those cell types. In MIC, 14 out of 19 colocalizing genes are colocalizing specifically in the MIC cell type, while 13 are also an eQTL another cell type, suggesting that the lack of overlap in other cell types is not due lack of statistical power. **c,** Region plots for the *BIN1* locus, showing colocalization specifically in MIC, while also being a significant eQTL in the EX, IN, and OLI cell types. **d,** Region plots for the *FERMT2* locus, showing colocalization specifically in MIC, while also being a significant eQTL in the EX and OLI cell types. Red dotted line indicates significance threshold for GWAS (p<5×10^-8^) or eQTL (q-value<0.05).

## Discussion

We present a comprehensive analysis of how genetic variation influences gene expression in specific cell types of the adult human brain. By leveraging snRNA-seq data from a large collection of samples and individuals, we were able to identify many *cis*-eQTLs, both shared and distinct to specific cell types. When comparing bulk and single-nucleus eQTLs between different tissues, we observed strong sharing of eQTLs between closely related cell types. Notably, sharing of eQTLs between cell types was substantially lower compared to bulk-derived eQTLs, suggesting that tissue-level averaging of cellular regulatory effects in bulk eQTLs might obscure cell-type-specific effects. By colocalizing eQTLs with GWAS hits, we identified high-confidence risk genes, their active cell types, and potential genetic variants for 11 brain related phenotypes.

We show that increasing the sample size enabled detection of 22,288 additional *cis*-eQTLs, which increased the cell-type-specificity compared to previous studies. We observed substantial agreement of eQTLs between closely related brain cell types, such as excitatory and inhibitory neurons. However, agreement was very low between more distinct cell types. Surprisingly, for some pairs of cell types, agreement between brain cell types was even lower than agreement of eQTLs between bulk brain and bulk blood. We subsequently compared the eQTLs identified per brain cell type to eQTLs identified using snRNA-seq in blood, where we observed highest consistency in findings from microglia and monocytes. These results suggest that the substantial agreement of eQTLs across different tissues that was previously reported^4,11,14^ is likely due to: (1) eQTLs for genes in cell programs that are shared between cells (e.g. housekeeping genes), (2) frequent but different cell types with similar biological functions that are shared between tissues (e.g. endothelial cells), (3) cell types derived from similar lineages (e.g. immune cells), or (4) averaging out of cell type specific effects in mixtures of cell types.

This study has several limitations. First, our resource consists of numerous datasets that used the same brain bank to collect their samples, most likely to balance age, gender, and disease status, or perhaps due to biopsy availability (**Supplementary Table 1**). Since our eQTL analysis uses linear regression, replicated samples across datasets would violate the underlying assumptions of the model; to address this, we included only one sample per individual. However, in the future, mixed-effects models should be considered to maximize the power within this resource^41–43^. Second, our efforts to quantify the cell-type-specificity of the *cis*-eQTLs do not explicitly account for the differences in statistical power in each of the cell types. The genes for which we have sufficient power in the low abundance cell types are often genes expressed in all cell types, possibly describing generic pathways active in all brain cell types. We observed that eQTLs for these low abundance cell types were more often shared with other cell types. Conversely, eQTLs in well powered cell types were more often distinct, possibly because these effects are cell-type-specific, or because there was insufficient power to detect them in other cell types. Until we have sufficient cells for all cell types, thereby minimizing the discrepancies in statistical power between them, we cannot conclude with complete confidence that an eQTL is cell-type-specific. Third, in our analysis we grouped cells based on the eight major cell types in the brain. However, there are many subtypes of these cells, and they have been shown to have specific eQTL effects^8^. We expect that future identification of cell-subtype eQTL effects will yield additional new findings. However, this will largely depend on the number of cells available for each of these subtypes. Lastly, our colocalization approach assumes a single causal variant, so we may have missed co-localizing signals in loci where this does not hold. This could be addressed in future by applying fine-mapping approaches^44,45^ to find independent signals and including this information in the colocalization step.

Our findings have significant implications for post-GWAS research, particularly for identifying causal genes and understanding the mechanisms underlying disease-associated loci. We prioritized 662 distinct genes for 11 brain-related traits and assigned putative driver cell types. Interestingly, we found that 68% of the genes colocalized in just one cell type. This suggests that disease risk at a particular GWAS locus is highly cell-type-specific. By demonstrating that eQTLs are highly cell-type-specific in the brain, we have highlighted the limitations of bulk eQTL studies for accurately linking genetic variants to their target genes. This suggests that many true eQTL associations may have been missed in previous analyses due to tissue-level averaging effects. By integrating single-cell eQTL data with GWAS findings, we can more precisely map genetic risk loci to their most relevant cell types, enabling better functional interpretation of disease-associated variants. As single-cell datasets continue to grow, expanding this approach to more tissues and cell types (e.g. single cell eQTLgen consortium^3^) will be critical for fully characterizing the regulatory landscape of genetic risk factors and improving the prioritization of therapeutic targets.

Overall, our study offers a comprehensive analysis of eQTLs across cell types in the human brain. We anticipate that this resource will prove highly valuable by pinpointing putative driver cell types behind disease-associated genetic variants. However, we expect that many additional eQTLs remain to be identified for low abundance yet highly disease-relevant cell types or subtypes. With the amount of single-nucleus data steadily increasing, we expect that resources like ours will grow quickly, enabling the interrogation of cell subtypes and cell programs at scale.

## Methods

### Data collection and description

We collected published single-nucleus brain RNA-seq samples from Synapse and the European Genome-Phenome Archive (EGA). An overview of the datasets and their accession IDs is provided in **Supplementary Table 1**. In total, we included 1,370 snRNA-seq samples and 1,210 genotype samples (non-unique, before quality control; QC) from the AMP-AD consortium (Mathys^18^, Sun^16^, Zhou^19^, Cain^20^, Fujita^8^, and Huang^21^), the PsychENCODE consortium (Ruzicka^22^) and Roche (RocheAD^7^, RocheMS^7^, and RocheColumbia^7^). 1,260 of the 1,370 samples could be linked to a genotype sample. Because of high overlap in genotype samples between the included datasets, this represented 785 unique individuals. For the creation of the final eQTL datasets, we selected the linked snRNA-seq sample that had the highest cell count for a given cell type and removed samples with too few cells (n<5). Consequentially, the final number of samples and individuals differs per cell type, and ranges between 649 and 757 unique individuals (**Supplementary Tables 1 and 2**).

### Analysis pipeline

For the processing of the data, we created an analysis pipeline building on previous work from the sc-eQTLgen consortium^3^, eQTLgen^4^, and Demuxafy^46^. This includes six separate analysis pipelines to reprocess raw RNA-seq data: (1) genotype imputation, (2) alignment and ambient RNA correction, (3) demultiplexing, doublet detection, and sample link validation, (4) cell type annotation, (5) eQTL mapping, and (6) a pipeline for the genotype data. The pipelines are set up to be performed in a federated manner, enabling easy inclusion of additional datasets without the need to share privacy-sensitive information. An initial version of these pipelines (except pipeline 2) were developed by the authors of their respective publications. In brief, changes included the addition of software required for the analysis of single-nucleus data (e.g. ambient RNA correction using CellBender^47^, doublet detection using scDblFinder^48^ and Doubletfinder^49^) and annotation of non-blood cell types. In addition, we made several improvements to the analysis workflow, including renormalization after sample QC and dynamic optimization of the number of expression PCs to include as covariates.

### Genotyping QC

The genotype data for the included datasets were generated using different platforms, including genotypes from whole-genome sequencing (WGS) (AMP-AD datasets) and genotyping arrays (Roche datasets and Ruzicka). For the other datasets, we imputed each dataset independently, except for the AMP-AD datasets, which were imputed collectively. For the WGS genotypes, we first split multiallelic sites into biallelic ones using bcftools^50^ norm (version 1.18). We removed variants with a Variant Quality Score Recalibration (VQSR) < 99.8 for SNPs and < 99.95 for indels. We then replaced uninformative genotype calls with missing (i.e. ‘./.’) using the following thresholds: read depth (DP) < 10, genotype quality (GQ) < 20, allelic balance (AB) > 0.2 for homozygous reference, AB < 0.8 for homozygous alternative and AB < 0.2 and > 0.8 for heterozygous. We then recalculated MAF and call rates (CR) and removed variants with a CR < 0.99, MAF < 0.01, or Hardy-Weinberg equilibrium p-value (HWE-p) < 1 x 10^-6^. For the genotype data based on arrays, we removed samples with missingness rates > 0.03 using plink2^51,52^ (version 20240205). If the input VCF was not on genome build GRCh38, we performed lift-over using CrossMap^53^ (version 0.6.6).

### Genotyping pre-processing

We then verified sex assignments using plink^51^ check-sex (version 20230116) and ancestry assignments using the 1000 Genomes GRCh38 30x^54^ reference (20220422 version) using plink2^51,52^. To maximize sample size, we imputed all ancestries together. First, we harmonized the strand alignment using GenotypeHarmonizer^55^ (version 1.4.27), fixed the strand orientation using bcftools^50^ fixref and norm, and removed variants with HWE-p < 1 x 10^-6^ and percentage missing > 0.05. We then removed samples with a heterozygosity rate > 3 standard deviations from the dataset average using vcftools^56^ het (version 0.1.16). To construct a kinship matrix, we used plink2^51,52^ -make-king, using only variants with a MAF > 0.05 and subsequently multiplying each KING-robust coefficient by two. We split the filtered VCF files per chromosome, phased the data using Eagle^57,58^ (version 2.4.1), and imputed using Minimac4^59^ (version 4.1.4) using the 1000G 30x reference^54^. After imputation, we removed variants with an imputation quality R^2^ < 0.3 and a MAF < 0.01, merged the individual chromosome files, and split the samples per sub-dataset.

### Single-nucleus RNA-seq analysis

All samples were processed with Cell Ranger (version 7.0.1) using the GRCh38 reference human genome by 10x (‘refdata-gex-GRCh38-2020-A’). We excluded 50 pools out of 165 from the Ruzicka dataset: two due to header mismatches and 48 due to low fraction of mapped genes. We performed ambient RNA correction by using CellBender^47^ (version 0.3.0) on the ‘raw_feature_bc_matrix’ Cell Ranger^60^ output, using default settings. Possible issues and warnings reported by CellBender were addressed according to the CellBender^47^ documentation. This twice resulted in halving the default learning-rate. We excluded three pools due to too few (< 100) unique molecular identifiers (UMIs) per cell. We then used the ambient RNA-adjusted gene expression quantifications for each nucleus from the CellBender output folder (‘cellbender_feature_bc_matrix_filtered’) for the following steps.

We identified doublets using both DoubletFinder^49^ (GitHub.com commit 1b1d4e2) and scDblFinder^48^ (version 1.8.0), using default settings. Barcodes identified as doublet by either method were marked as doublets. If the RNA-seq data was multiplexed, we applied souporcell^20^ (version 2.5) to identify doublets and assign droplets to individuals. To improve scalability, we created a custom snakemake^61^ version of the souporcell pipeline that enables pause-and-resume and parallel processing of individual steps on an HPC environment. We ran souporcell^62^ using skip_remap, but with otherwise default settings. If the RNA-seq was not multiplexed, we applied VerifyBamID^63^ (version 1.1.3) to identify sample swaps. Samples with a sequence and array estimate of contamination (CHIPMIX) > 0.02 were manually inspected for mix-ups. If the CHIPMIX was approximately 0.0 and CHIP_ID matched the genotype assignment, we kept the original assignment. If CHIPMIX was approximately 1.0, we discarded the genotype assignment. In both the demultiplexing and sample swap detection, we first filtered the Cell Ranger^60^ BAM file (‘possorted_genome_bam’) to keep exonic regions because this improved computational efficiency without affecting the individual assignments.

Simultaneously, we annotated each cell using label transfer with Azimuth^64^. We used the default human motor cortex single-nucleus reference from the Allen Institute^65^ as reference. In brief, this dataset consists of single-nucleus data from two human individuals annotated at different subclasses of brain cell types: astrocyte (AST), endothelial cells (END), eight glutamatergic neuron subtypes (EX), six GABAergic neuron subtypes (IN), perivascular macrophage (MIC), oligodendrocyte (OLI), oligodendrocyte precursor cell (OPC), and brain pericyte (PER). Per sequencing run, we took the CellBender-corrected gene expression quantifications and applied SCTransform, FindTransferAnchors (dims = 50), and MapQuery using Seurat^64^ (version 4.4.0). For downstream analyses, we merged subtypes of neurons into their overarching excitatory glutamatergic (EX) and inhibitory GABAergic (IN) cell types, reducing the total number of distinct cell types to eight.

We then performed barcode QC to select high quality cells. Here, we observed that the number of nuclei per pool far exceeded experimental expectations due to the use of CellBender. To combat this, we capped the number of nuclei per pool to the average number of nuclei according to CellRanger over all pools + 20%. We added the 20% to account for additional power to detect cells due to ambient removal. This threshold was determined per dataset and was 8,471 for RocheAD, 6,973 for RocheMS, 9,350 for RocheColumbia, 10,782 for Cain, 1,762 for Mathys, 5,086 for Zhou, 23,626 for Fujita, 6,760 for Sun, 4,978 for Huang, and 36,319 for Ruzicka. If a pool exceeded this maximum, we included only the nuclei with the highest number of UMIs up to the maximum amount. Furthermore, nuclei were defined as barcodes with at least 500 UMIs and < 5% mitochondrial RNA that were marked as singlet by both doublet-detection approaches. Finally, samples with < 5 cells were excluded.

### eQTL mapping

For the eQTL analysis, we first excluded cells that were not assigned to a genotype sample. Individuals with multiple samples within one dataset were combined. For individuals with samples in multiple datasets, we only included the sample with the highest number of cells. We then generated pseudo-bulk gene expression matrices for each cell type by summing all counts for each gene in each individual. Next, we removed genes that showed zero variability, had fewer than 10 individuals with at least one read, or had an average counts per million (CPM) < 1 over all samples. The resulting gene by sample matrix was then normalized using trimmed mean of M values (TMM) using edgeR^66^, after which we applied principal component analysis (PCA) to summarize the expression matrix. We excluded 10−20 samples (dependent on the cell type; **Supplementary Table 2**) that had an absolute loading z-score > 3 on PC1−3 and reperformed the PCA on the remaining samples. We then aimed to correct the gene expression data for the resulting PCs so that 80% of the gene expression variance was removed. We based this selection criterium on previous work by Bryois *et al.*^7^, who suggested this was the optimum for single-nucleus eQTL analysis. We determined that 80% of the variance was captured by 98 expression PCs in the EX cell type and used the same cutoff for the remaining cell types. Per cell type, we additionally corrected the gene expression for dataset indicator variables and the 98 expression PCs using ordinary least squares (OLS) regression. We then mapped *cis*-eQTLs within a 1-Mb window of the TSS of each gene using mbQTL (version 1.5.1)^14^ with Pearson correlations (--norank). Only variants with > 9 observations, MAF > 0.05, HWE-p > 0.0001, and call rate > 0.95 were considered. We also required variants to have at least two observations in each of three genotype groups. We performed 1,000 permutations per gene to determine an empirically adjusted gene-level p-value using the beta distribution estimation approach^14,67^. Per gene, the lead variant was selected based on the lowest nominal p-value, and multiple testing correction across all genes (per cell type) was performed using the q-value^68^ R package. Genes with a q-value<0.05 were considered significant.

### eQTL replication

We evaluated the replication of eQTL effects based on three different measurements of agreement:AC, *π*_1_^69^, and *R_b_*^15^. In brief, AC describes the proportion of effects with a shared direction, *π*_1_ estimates the proportion of true positives in the replication dataset, and *R_b_* estimates the correlation of effect direction. For more information, see de Klein *et al.*^14^. To calculate this, we took the lead variant per gene that was significant (q-value<0.05) in the discovery dataset and compared this with the exact same variant-gene in the replication dataset. This means that an eQTL agreement is not necessarily significant in the replication dataset when calculating the AC, *π_1_*, or *R_b_*. To determine how many effects are replicating significantly, we calculated a BH-FDR using the nominal p-values from the replication dataset for overlapping eQTLs.

### Single nucleus PBMC replication dataset

PBMCs were collected from 123 donors from the general population and applied to the 10X Genomics Multiome^70^ v3 sequencing platform. In parallel, we performed genome-wide genotyping using the Illumina GSA-MD v3 array, followed by imputation using the 1000G 30x reference^54^, yielding a total of 50,998,911 genetic variants. As quality control of the single nucleus data, first, doublets arising from different individuals were identified using Souporcell^62^, which clusters cells based on genetic variants present in the 3-prime end of the transcripts. This revealed that, on average, 27.1% of the captured cells were doublets, which were subsequently excluded from further analysis. The genotypes of the Souporcell clusters were subsequently correlated to GSA genotypes, allowing cells to be assigned to samples. Additional stringent quality control was applied to retain only high-quality cells that had paired RNA and ATAC data, resulting in a final dataset of 224,202 cells suitable for downstream analysis.

For data integration, we employed a weighted nearest-neighbour (WNN) approach to jointly embed scRNA-seq and scATAC-seq modalities. Cell type annotation was then performed using the Azimuth reference framework, using the publicly available 10X Multiome dataset of 10,000 PBMCs from a single donor that is available in the SeuratData R package^71^. This allowed us to identify eight major cell types (’B’, ’CD4+ T’, ’CD8+ T ’, ’DC’, ’monocyte’, ’NK’, ’plasmablast’ and ’erythrocytes’). The single-nucleus expression data was first normalized by calculating the total UMI per nucleus, then dividing total UMIs per nucleus by this mean to get a scaling factor, then applying this scaling factor to all gene counts for each nucleus (i.e. PFlog1pPF^72^ normalization). Expression was next pseudobulked by summing the expression for each combination of cell type and sample. The first 10 principal components were then calculated over these pseudobulks for each cell type separately. We then fit a linear mixed model using the sc-eQTLgen QTL mapping tool LIMIX-QTL^43,73^, predicting gene expression by genetic variants, while correcting for the first 10 expression PCs as fixed effects, and the donor assignments of the samples as random effects. We confined our analysis to *cis*-eQTL effects within a windows size of 1-Mbfrom the gene body. We used 20 variant-level permutations for each gene to control false discovery rate at the variant level and then performed q-value correction^68^ over the strongest permutation-level effects per gene to account for gene-level false discovery. We considered genes with a gene-level q-value of < 0.05 as significant associations.

### Colocalization

Using the Coloc^74^ package (v5.2.3) and the GWAS summary statistics for AD^24^, ADHD^75^, ALS^27^, BPD^31^, epilepsy^32^, Intelligence^76^, MDD^29^, MS^25^, PD^26^ , SCZ^77^ and years of schooling^34^, we conducted a colocalization analysis to identify potential shared genetic signals between the disease/trait and gene expression. First, we identified significant variants (p<5 x 10^-8^) in the GWAS summary statistics. We then selected genes with a TSS within 500 kb (upstream and downstream) of these variants. We confined ourselves to genes with an eQTL q-value<0.05. For each selected gene, we extracted all variants located within 500 kb of the TSS (upstream and downstream) from the GWAS summary statistics and all overlapping variants from the eQTL data. Finally, we performed colocalization analysis with the coloc.abf() function from the Coloc^74^ package, using the default priors. This function returns posterior probabilities for five hypotheses: PPH0 (no association with eQTL or GWAS), PPH1 (association with eQTL but not with GWAS), PPH2 (association with GWAS but not with eQTL), PPH3 (association with both eQTL and GWAS, but no shared causal variant), and PPH4 (association with both eQTL and GWAS with a shared causal variant). A locus was considered colocalizing if the posterior probability for PPH4 was > 0.8. On top of that, we required that at least one of the overlapping variants in the locus had a GWAS p-value<5×10^-8^, and a significant eQTL p-value. Finally, we used LocusBreaker^35^ and determined independent loci for each GWAS to determine how many loci had at least one colocalizing gene. We note that since we performed the LocusBreaker analysis after the colocalization analysis and because the method does not use linkage disequilibrium information, some colocalizing genes do not overlap LocusBreaker loci and numbers of loci for a given GWAS may differ from their original publications.

## Ethical compliance

All cohorts included in this study enrolled participants with informed consent and collected and analyzed data in accordance with ethical and institutional regulations. The information about individual institutional review board approvals is available in the original publications for each cohort (see ‘Data availability’). Where applicable, data access agreements were signed by the investigators before acquisition of the data, and these state the data usage terms. To protect the privacy of the participants, data access was restricted to the investigators of this study, as defined in those data access agreements. Following the data use agreements, only summarylZJlevel data is made publicly available, and it is strictly mentioned in the disclaimer that data cannot be used to relZJidentify study participants.

The protocol of the scMetaBrain study was reviewed by the Medical Ethics Review Board (CTc) of the University Medical Center Groningen under research registry number 16017, which concluded that it is not a clinical research study with human subjects as meant in the Dutch Medical Research Involving Human Subjects Act (WMO). PBMC snRNA-seq dataset: The LifeLines DEEP study was approved by the ethics committee of the University Medical Centre Groningen, document number METC UMCG LLDEEP: M12.113965. All participants signed informed consent from prior to study enrollment. All procedures performed in studies involving human participants were in accordance with the ethical standards of the institutional and/or national research committee and with the 1964 Helsinki declaration and its later amendments or comparable ethical standards.

## Data availability

Summary statistics for the *cis-*eQTL analyses we performed will be made available after publication. PBMC snRNA-seq data for 123 donors will be made available upon publication. Our study is comprised of previously published human brain eQTL datasets. All of these datasets are available on request or through online repositories after signing a data access agreement. The mode of access for each of the included datasets was as follows. RocheAD^7^ (accession code: <u>EGAD00001009166</u>), RocheMS^7^ (accession code: <u>EGAD00001009169</u>), and RocheColumbia^7^ (accession code: <u>EGAD00001009168</u>) data was downloaded from EGA using <u>pyEGA3</u> download client. ROSMAP (accession code: <u>syn3219045</u>), Cain^20^ and Fujita^8^ (accession code: <u>syn23650894</u>), Mathys^18^ (accession code: <u>syn18485175</u>), Zhou^19^ (accession code: <u>syn21670836</u>), Sun^16^ (accession code: <u>syn52293417</u>), Huang^21^ (accession code: <u>syn50670858</u>), and Ruzicka^22^ (accession code: <u>syn22755055</u>) data were downloaded from Synapse.org using synapseclient (version 3.2.0). The Bryois *et al.* (accession code: <u>10.5281/zenodo.7276971</u>) and Fujita *et al.* eQTL summary statistics were downloaded from Zenodo (accession code <u>syn52335732</u>). The MetaBrain eQTL summary statistics were downloaded from metabrain.nl. GTEx-v8 summary statistics were downloaded from the GTEx portal website^2^. Source data are provided with this paper. GWAS data was downloaded from GWAS catalog (AD, ALS, MDD, PD, intelligence), PGC (SCZ, ADHD, BPD), and Open GWAS Project (years of schooling).

## Acknowledgments

We thank the donors of the brain tissues underlying the RNA-seq data used for this study and their families for their willingness to donate samples for research. We also thank all the researchers involved with the included cohorts for making their data available for use. We thank the UMCG Genomics Coordination Center, the UG Center for Information Technology, and their sponsors BBMRI-NL and TarGet for storage and computing infrastructure. The authors thank Drew Neavin, Urmo Võsa, Robert Warmerdam, Jose Alquicira Hernandez, and Lieke Michielsen for their support in using their software. We also thank Kate Mc Intyre for the editorial assistance.

## AMP-AD

The results published here are in whole or in part based on data obtained from the AD Knowledge Portal.

## The Religious Orders Study and Memory and Aging Project (ROSMAP) Study

Study data were provided by the Rush Alzheimer’s Disease Center, Rush University Medical Center, Chicago. Data collection was supported through funding by NIA grants RF1AG57473 (single nucleus RNAseq, U01AG61356 (whole genome sequencing, targeted proteomics, ROSMAP AMP-AD), the Illinois Department of Public Health (ROSMAP), and the Translational Genomics Research Institute (genomic). Study data were generated from postmortem brain tissue provided by the Religious Orders Study and Rush Memory and Aging Project (ROSMAP) cohort at Rush Alzheimer’s Disease Center, Rush University Medical Center, Chicago. This work was funded by NIH grants U01AG061356 (De Jager/Bennett), RF1AG057473 (De Jager/Bennett), and U01AG046152 (De Jager/Bennett) as part of the AMP-AD consortium, as well as NIH grants R01AG066831 (Menon) and U01AG072572 (De Jager/St George-Hyslop).

## The MIT ROSMAP Single-Nucleus Multiomics Study (MITROSMAPMultiomics)

Study data were generated from postmortem brain tissue provided by the Religious Orders Study and Rush Memory and Aging Project (ROSMAP) cohort at Rush Alzheimer’s Disease Center, Rush University Medical Center, Chicago. This work was supported in part by the Cure Alzheimer’s Fund, NIH grants AG058002, AG062377, NS110453, NS115064, AG062335, AG074003, NS127187, MH119509, HG008155 (M.K.), RF1AG062377, RF1 AG054321, RO1 AG054012 (L.-H.T.) and the NIH training grant GM087237 (to C.A.B.). ROSMAP is supported by P30AG10161, P30AG72975, R01AG15819, R01AG17917. U01AG46152, U01AG61356.

## The ROSMAP Mammillary Body Study (ROSMAP_MammillaryBody)

Study data were generated from postmortem brain tissue provided by the Religious Orders Study and Rush Memory and Aging Project (ROSMAP) cohort at Rush Alzheimer’s Disease Center, Rush University Medical Center, Chicago. This research was supported in part by JBP foundation, Ludwig family foundation, and National Institute of Health (NIH) grants RF1AG054321, RF1AG062377, RF1AG054012, U01NS110453, R01AG062335, and R01AG058002 (L-HT); P30AG10161, R01AG15819, and R01AG1717 (DAB); RF1AG054012, U01NS110453, R01AG062335, and R01AG058002 (MK).

## PsychENCODE

Data were generated as part of the PsychENCODE Consortium. Visit 10.7303/syn26365932 for a complete list of grants and PIs. Data were generated as part of the PsychENCODE Consortium, supported by: U01DA048279, U01MH103339, U01MH103340, U01MH103346, U01MH103365, U01MH103392, U01MH116438, U01MH116441, U01MH116442, U01MH116488, U01MH116489, U01MH116492, U01MH122590, U01MH122591, U01MH122592, U01MH122849, U01MH122678, U01MH122681, U01MH116487, U01MH122509, R01MH094714, R01MH105472, R01MH105898, R01MH109677, R01MH109715, R01MH110905, R01MH110920, R01MH110921, R01MH110926, R01MH110927, R01MH110928, R01MH111721, R01MH117291, R01MH117292, R01MH117293, R21MH102791, R21MH103877, R21MH105853, R21MH105881, R21MH109956, R56MH114899, R56MH114901, R56MH114911, R01MH125516, and P50MH106934 awarded to: Alexej Abyzov, Nadav Ahituv, Schahram Akbarian, Alexander Arguello, Lora Bingaman, Kristin Brennand, Andrew Chess, Gregory Cooper, Gregory Crawford, Stella Dracheva, Peggy Farnham, Mark Gerstein, Daniel Geschwind, Fernando Goes, Vahram Haroutunian, Thomas M. Hyde, Andrew Jaffe, Peng Jin, Manolis Kellis, Joel Kleinman, James A. Knowles, Arnold Kriegstein, Chunyu Liu, Keri Martinowich, Eran Mukamel, Richard Myers, Charles Nemeroff, Mette Peters, Dalila Pinto, Katherine Pollard, Kerry Ressler, Panos Roussos, Stephan Sanders, Nenad Sestan, Pamela Sklar, Nick Sokol, Matthew State, Jason Stein, Patrick Sullivan, Flora Vaccarino, Stephen Warren, Daniel Weinberger, Sherman Weissman, Zhiping Weng, Kevin White, A. Jeremy Willsey, Hyejung Won, and Peter Zandi.

## Funding

J.D. is an employee of Roche/Genentech. Y.H. and E.A.T. are employees of Biogen. L.F. is supported by a grant from the Dutch Research Council (ZonMWlZJVICI 09150182010019), an Oncode Senior Investigator grant, and a sponsored research collaboration with Biogen and Roche. M.W. is supported by a grant from the Dutch Research Council (NWO Vidi 223.041).

## Author contributions

**Martijn Vochteloo**: Conceptualization (supporting), Data curation (lead), Formal Analysis (lead), Investigation (lead), Methodology (equal), Project Administration (equal), Software (lead), Validation (lead), Visualization (lead), Writing - Original Draft (lead), Writing - Review & Editing (supporting). **Anoek Kooijmans:** Formal Analysis (equal), Methodology (supporting), Software (equal), Validation (supporting), Visualization (supporting). **Joost Bakker:** Formal Analysis (supporting), Software (supporting), Visualization (supporting). **Roy Oelen**: Software (supporting). **Jelmer Niewold:** Data generation. **Dan Kaptijn**: Software (supporting). **Marc-Jan Bonder**: Software (supporting), Supervision (supporting). **Monique van der Wijst**: Supervision (supporting). **Yunfeng Huang:** Conceptualization (lead), Funding Acquisition (equal), Writing - Review & Editing (supporting). **Dennis Baird**: Conceptualization (supporting). **Julien Bryois:** Conceptualization (lead), Funding Acquisition (lead), Writing - Review & Editing (supporting). **Ellen Tsai:** Conceptualization (lead), Funding Acquisition (lead), Writing - Review & Editing (supporting). **Lude Franke:** Conceptualization (lead), Funding Acquisition (lead), Methodology (lead), Project Administration (equal), Resources (lead), Supervision (equal), Visualization (supporting), Writing - Review & Editing (supporting). **Harm-Jan Westra:** Conceptualization (lead), Data curation (supporting), Formal Analysis (supporting), Funding Acquisition (equal), Investigation (equal), Methodology (lead), Project Administration (lead), Software (supporting), Supervision (lead), Visualization (lead), Writing - Review & Editing (lead). Roles as defined by: CRediT (Contributor Roles Taxonomy).

## Competing interests

J.B. is an employee of Roche/Genentech. Y.H. and E.A.T. are employees of Biogen. All other authors declare no competing interests.

## Supplementary Figures

**Supplementary Figure 1 | Replication statistics comparing scMetaBrain to bulk and single cell datasets.** eQTLs from specific cell types show best replication statistics (R_b_) when compared to eQTLs from single cell and bulk tissues.

**Supplementary Figure 2 | Colocalization of AD and PD GWAS in GRN locus.** Multiple genes located in the GRN locus show colocalization with PD, but not AD. Non-coding genes were excluded from this figure. Red lines indicate multiple testing correction threshold for GWAS (p<5×10^-8^) or eQTL (q-value<0.05).

**Supplementary Figure 3 | Genes that show colocalization with AD specifically in MIC.** 13 genes showed colocalization with AD specifically in MIC, while many of them were also significant eQTLs in other cell types. Non-coding genes were excluded from this figure. Red lines indicate multiple testing correction threshold for GWAS (p<5×10^-8^) or eQTL (q-value<0.05).

**Supplementary Tables**

## Supplementary Tables

Supplementary Table 1: Dataset overview.

Supplementary Table 2: eQTL QC overview.

Supplementary Table 3: eQTL results per cell type.

Supplementary Table 4. eQTL agreement between cell types and tissues.

Supplementary Table 5: GWAS colocalization results.

Supplementary Table 6: Summary of GWAS colocalization results

## References

1. E, S., et al. The NHGRI-EBI GWAS Catalog: knowledgebase and deposition resource. Nucleic Acids Res. 51, (2023).

2. The GTEx Consortium et al. The GTEx Consortium atlas of genetic regulatory effects across human tissues. Science 369, 1318–1330 (2020).

3. van der Wijst, M. et al. The single-cell eQTLGen consortium. eLife 9, e52155 (2020).

4. Võsa, U. et al. Large-scale cis- and trans-eQTL analyses identify thousands of genetic loci and polygenic scores that regulate blood gene expression. Nat. Genet. 53, 1300–1310 (2021).

5. Wang, D. et al. Comprehensive functional genomic resource and integrative model for the human brain. Science 362, eaat8464 (2018).

6. Ng, B. et al. An xQTL map integrates the genetic architecture of the human brain’s transcriptome and epigenome. Nat. Neurosci. 20, 1418–1426 (2017).

7. Bryois, J. et al. Cell-type-specific cis-eQTLs in eight human brain cell types identify novel risk genes for psychiatric and neurological disorders. Nat. Neurosci. 25, 1104–1112 (2022).

8. Fujita, M. et al. Cell subtype-specific effects of genetic variation in the Alzheimer’s disease brain. Nat. Genet. https://doi.org/10.1038/s41588-024-01685-y (2024) doi:10.1038/s41588-024-01685-y.

9. Lonsdale, J. et al. The Genotype-Tissue Expression (GTEx) project. Nat. Genet. 45, 580–585 (2013).

10. The GTEx Consortium et al. The Genotype-Tissue Expression (GTEx) pilot analysis: Multitissue gene regulation in humans. Science 348, 648–660 (2015).

11. GTEx Consortium. Genetic effects on gene expression across human tissues. Nature 550, 204–213 (2017).

12. Donovan, M. K. R., D’Antonio-Chronowska, A., D’Antonio, M. & Frazer, K. A. Cellular deconvolution of GTEx tissues powers discovery of disease and cell-type associated regulatory variants. Nat. Commun. 11, 955 (2020).

13. Raj, T. et al. Integrative transcriptome analyses of the aging brain implicate altered splicing in Alzheimer’s disease susceptibility. Nat. Genet. 50, 1584–1592 (2018).

14. De Klein, N. et al. Brain expression quantitative trait locus and network analyses reveal downstream effects and putative drivers for brain-related diseases. Nat. Genet. 55, 377–388 (2023).

15. Qi, T. Identifying gene targets for brain-related traits using transcriptomic and methylomic data from blood. Nat. Commun. 9, 2282 (2018).

16. Sun, N. et al. Single-nucleus multiregion transcriptomic analysis of brain vasculature in Alzheimer’s disease. Nat. Neurosci. https://doi.org/10.1038/s41593-023-01334-3 (2023) doi:10.1038/s41593-023-01334-3.

17. Zeng, B. et al. Genetic regulation of cell type–specific chromatin accessibility shapes brain disease etiology. Science 384, eadh4265 (2024).

18. Mathys, H. et al. Single-cell transcriptomic analysis of Alzheimer’s disease. Nature 570, 332–337 (2019).

19. Zhou, Y. et al. Human and mouse single-nucleus transcriptomics reveal TREM2-dependent and TREM2-independent cellular responses in Alzheimer’s disease. Nat. Med. 26, 131–142 (2020).

20. Cain, A. et al. Multicellular communities are perturbed in the aging human brain and Alzheimer’s disease. Nat. Neurosci. 26, 1267–1280 (2023).

21. Huang, W.-C. et al. Lateral mammillary body neurons in mouse brain are disproportionately vulnerable in Alzheimer’s disease. Sci. Transl. Med. 15, eabq1019 (2023).

22. Ruzicka, W. B., et al. Single-Cell Dissection of Schizophrenia Reveals Neurodevelopmental-Synaptic Axis and Transcriptional Resilience. http://medrxiv.org/lookup/doi/10.1101/2020.11.06.20225342 (2020) doi:10.1101/2020.11.06.20225342.

23. Karczewski, K. J. et al. The mutational constraint spectrum quantified from variation in 141,456 humans. Nature 581, 434–443 (2020).

24. Bellenguez, C. et al. New insights into the genetic etiology of Alzheimer’s disease and related dementias. Nat. Genet. 54, 412–436 (2022).

25. International Multiple Sclerosis Genetics Consortium et al. Multiple sclerosis genomic map implicates peripheral immune cells and microglia in susceptibility. Science 365, eaav7188 (2019).

26. Program (GP2), T. G. P. G. & Leonard, H. L. Novel Parkinson’s Disease Genetic Risk Factors Within and Across European Populations. 2025.03.14.24319455 Preprint at 10.1101/2025.03.14.24319455 (2025).

27. Van Rheenen, W. et al. Common and rare variant association analyses in amyotrophic lateral sclerosis identify 15 risk loci with distinct genetic architectures and neuron-specific biology. Nat. Genet. 53, 1636–1648 (2021).

28. Trubetskoy, V. et al. Mapping genomic loci implicates genes and synaptic biology in schizophrenia. Nature 604, 502–508 (2022).

29. Adams, M. J. et al. Trans-ancestry genome-wide study of depression identifies 697 associations implicating cell types and pharmacotherapies. Cell 188, 640–652.e9 (2025).

30. Demontis, D. et al. Genome-wide analyses of ADHD identify 27 risk loci, refine the genetic architecture and implicate several cognitive domains. Nat. Genet. 55, 198–208 (2023).

31. O’Connell, K. S. et al. Genomics yields biological and phenotypic insights into bipolar disorder. Nature 639, 968–975 (2025).

32. The International League Against Epilepsy Consortium on Complex Epilepsies et al. Genome-wide mega-analysis identifies 16 loci and highlights diverse biological mechanisms in the common epilepsies. Nat. Commun. 9, 5269 (2018).

33. Davies, G. et al. Study of 300,486 individuals identifies 148 independent genetic loci influencing general cognitive function. Nat. Commun. 9, 2098 (2018).

34. 23andMe Research Team et al. Gene discovery and polygenic prediction from a genome-wide association study of educational attainment in 1.1 million individuals. Nat. Genet. 50, 1112–1121 (2018).

35. Pirastu, N. et al. GWAS for male-pattern baldness identifies 71 susceptibility loci explaining 38% of the risk. Nat. Commun. 8, 1584 (2017).

36. Jian, J., Konopka, J. & Liu, C. Insights into the role of progranulin in immunity, infection, and inflammation. J. Leukoc. Biol. 93, 199–208 (2013).

37. Nalls, M. A. et al. Evidence for GRN connecting multiple neurodegenerative diseases. Brain Commun. 3, fcab095 (2021).

38. Sudwarts, A. et al. BIN1 is a key regulator of proinflammatory and neurodegeneration-related activation in microglia. Mol. Neurodegener. 17, 33 (2022).

39. Taga, M. et al. BIN1 protein isoforms are differentially expressed in astrocytes, neurons, and microglia: neuronal and astrocyte BIN1 are implicated in tau pathology. Mol. Neurodegener. 15, 44 (2020).

40. Chapuis, J. et al. Genome-wide, high-content siRNA screening identifies the Alzheimer’s genetic risk factor FERMT2 as a major modulator of APP metabolism. Acta Neuropathol. (Berl.) 133, 955–966 (2017).

41. Davenport, E. E. et al. Discovering in vivo cytokine-eQTL interactions from a lupus clinical trial. Genome Biol. 19, 168 (2018).

42. Zeng, B. et al. Multi-ancestry eQTL meta-analysis of human brain identifies candidate causal variants for brain-related traits. Nat. Genet. 54, 161–169 (2022).

43. Lippert, C., Casale, F. P., Rakitsch, B. & Stegle, O. LIMIX: genetic analysis of multiple traits. bioRxiv 003905 (2014) doi:10.1101/003905.

44. Wallace, C. A more accurate method for colocalisation analysis allowing for multiple causal variants. PLOS Genet. 17, e1009440 (2021).

45. Wang, G., Sarkar, A., Carbonetto, P. & Stephens, M. A simple new approach to variable selection in regression, with application to genetic fine mapping. J. R. Stat. Soc. Ser. B Stat. Methodol. 82, 1273–1300 (2020).

46. Neavin, D. et al. Demuxafy: improvement in droplet assignment by integrating multiple single-cell demultiplexing and doublet detection methods. Genome Biol. 25, 94 (2024).

47. Fleming, S. J. et al. Unsupervised removal of systematic background noise from droplet-based single-cell experiments using CellBender. Nat. Methods https://doi.org/10.1038/s41592-023-01943-7 (2023) doi:10.1038/s41592-023-01943-7.

48. Germain, P.-L., Lun, A., Meixide, C. G., Macnair, W. & Robinson, M. D. Doublet identification in single-cell sequencing data. F1000research (2022).

49. McGinnis, C. S., Murrow, L. M. & Gartner, Z. J. DoubletFinder: Doublet Detection in Single-Cell RNA Sequencing Data Using Artificial Nearest Neighbors. Cell Syst. 8, 329–337.e4 (2019).

50. Danecek, P. et al. Twelve years of SAMtools and BCFtools. GigaScience 10, giab008 (2021).

51. Purcell, S. et al. PLINK: a tool set for whole-genome association and population-based linkage analyses. Am. J. Hum. Genet. 81, 559–575 (2007).

52. Chang, C. C. et al. Second-generation PLINK: rising to the challenge of larger and richer datasets. Gigascience 4, s13742-015-0047–8 (2015).

53. Zhao, H. et al. CrossMap: a versatile tool for coordinate conversion between genome assemblies. Bioinformatics 30, 1006–1007 (2014).

54. Byrska-Bishop, M. et al. High-coverage whole-genome sequencing of the expanded 1000 Genomes Project cohort including 602 trios. Cell 185, 3426–3440.e19 (2022).

55. Deelen, P. et al. Genotype harmonizer: automatic strand alignment and format conversion for genotype data integration. BMC Res. Notes 7, 901 (2014).

56. Danecek, P. et al. The variant call format and VCFtools. Bioinformatics 27, 2156–2158 (2011).

57. Loh, P.-R., Palamara, P. F. & Price, A. L. Fast and accurate long-range phasing in a UK Biobank cohort. Nat. Genet. 48, 811–816 (2016).

58. Loh, P.-R. et al. Reference-based phasing using the Haplotype Reference Consortium panel. Nat. Genet. 48, 1443–1448 (2016).

59. Das, S. et al. Next-generation genotype imputation service and methods. Nat. Genet. 48, 1284–1287 (2016).

60. Zheng, G. X. Y. et al. Massively parallel digital transcriptional profiling of single cells. Nat. Commun. 8, 14049 (2017).

61. Mölder, F. et al. Sustainable data analysis with Snakemake. F1000Research 10, 33 (2021).

62. Heaton, H. et al. Souporcell: robust clustering of single-cell RNA-seq data by genotype without reference genotypes. Nat. Methods 17, 615–620 (2020).

63. Jun, G. et al. Detecting and Estimating Contamination of Human DNA Samples in Sequencing and Array-Based Genotype Data. Am. J. Hum. Genet. 91, 839–848 (2012).

64. Hao, Y. et al. Integrated analysis of multimodal single-cell data. Cell 184, 3573–3587.e29 (2021).

65. Bakken, T. E. et al. Comparative cellular analysis of motor cortex in human, marmoset and mouse. Nature 598, 111–119 (2021).

66. Chen, Y., Chen, L., Lun, A. T. L., Baldoni, P. L. & Smyth, G. K. edgeR v4: powerful differential analysis of sequencing data with expanded functionality and improved support for small counts and larger datasets. Nucleic Acids Res. 53, gkaf018 (2025).

67. Ongen, H., Buil, A., Brown, A. A., Dermitzakis, E. T. & Delaneau, O. Fast and efficient QTL mapper for thousands of molecular phenotypes. Bioinformatics 32, 1479–1485 (2016).

68. Storey, J., Bass, A., Dabney, A. & Robinson, D. qvalue: Q-value estimation for false discovery rate control. (2022).

69. Holland, D. et al. Estimating Effect Sizes and Expected Replication Probabilities from GWAS Summary Statistics. Front. Genet. 7, (2016).

70. Conklin, K., et al. 10X Genomics Single-Nucleus Multiome (RNA + ATAC) Assay for Profiling Adult Human Tissues. https://protocols.io/view/10x-genomics-single-nucleus-multiome-rna-atac-assa-b93fr8jn (2022).

71. Hao, Y. et al. Dictionary learning for integrative, multimodal and scalable single-cell analysis. Nat. Biotechnol. 42, 293–304 (2024).

72. Booeshaghi, A. S., Hallgrímsdóttir, I. B., Gálvez-Merchán, Á. & Pachter, L. Depth normalization for single-cell genomics count data. 2022.05.06.490859 Preprint at 10.1101/2022.05.06.490859 (2022).

73. Cuomo, A. S. E. et al. Optimizing expression quantitative trait locus mapping workflows for single-cell studies. Genome Biol. 22, 188 (2021).

74. Giambartolomei, C. et al. Bayesian Test for Colocalisation between Pairs of Genetic Association Studies Using Summary Statistics. PLoS Genet. 10, e1004383 (2014).

75. Demontis, D. et al. Genome-wide analyses of ADHD identify 27 risk loci, refine the genetic architecture and implicate several cognitive domains. Nat. Genet. 55, 198–208 (2023).

76. Savage, J. E. et al. Genome-wide association meta-analysis in 269,867 individuals identifies new genetic and functional links to intelligence. Nat. Genet. 50, 912–919 (2018).

77. Howard, D. M. et al. Genome-wide association study of depression phenotypes in UK Biobank identifies variants in excitatory synaptic pathways. Nat. Commun. 9, 1470 (2018).

