## Supplementary figures and images for "High cell-type specificity of eQTLs revealed by single-nucleus analyses of brain and blood"

### Supplementary Figure 1

#### Supplementary Figure 1

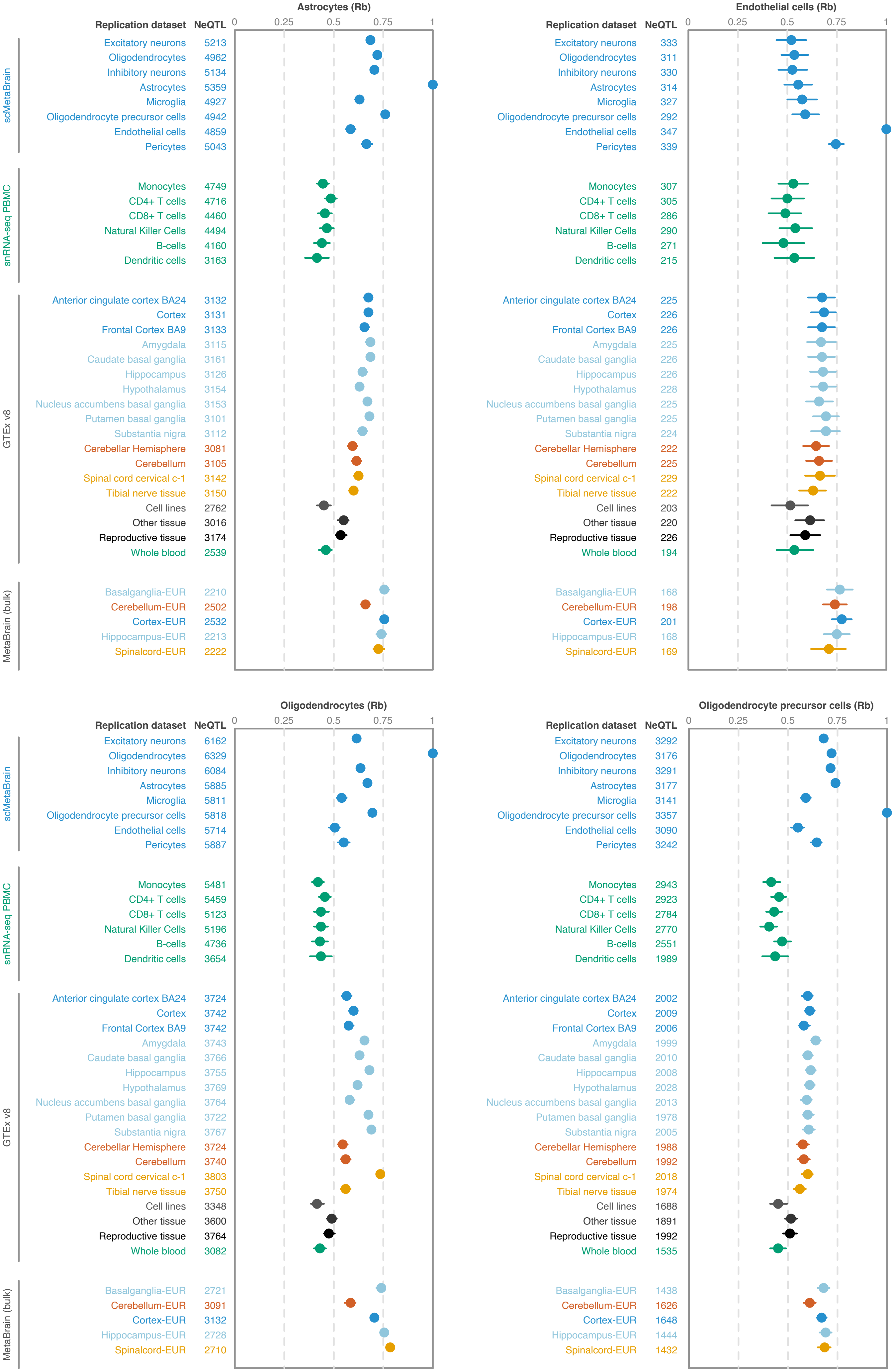

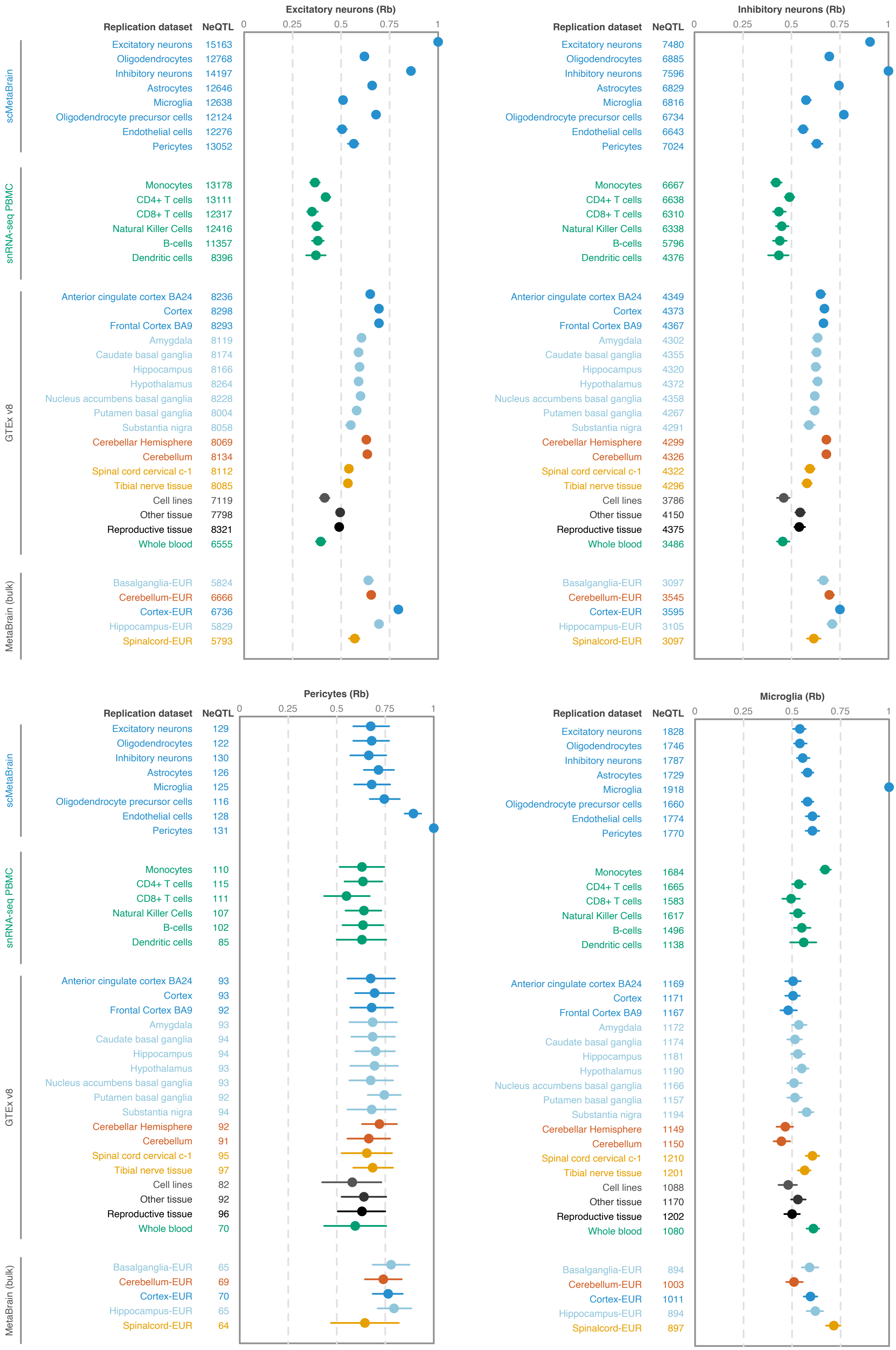

### Supplementary Figure 2

#### Supplementary Figure 2

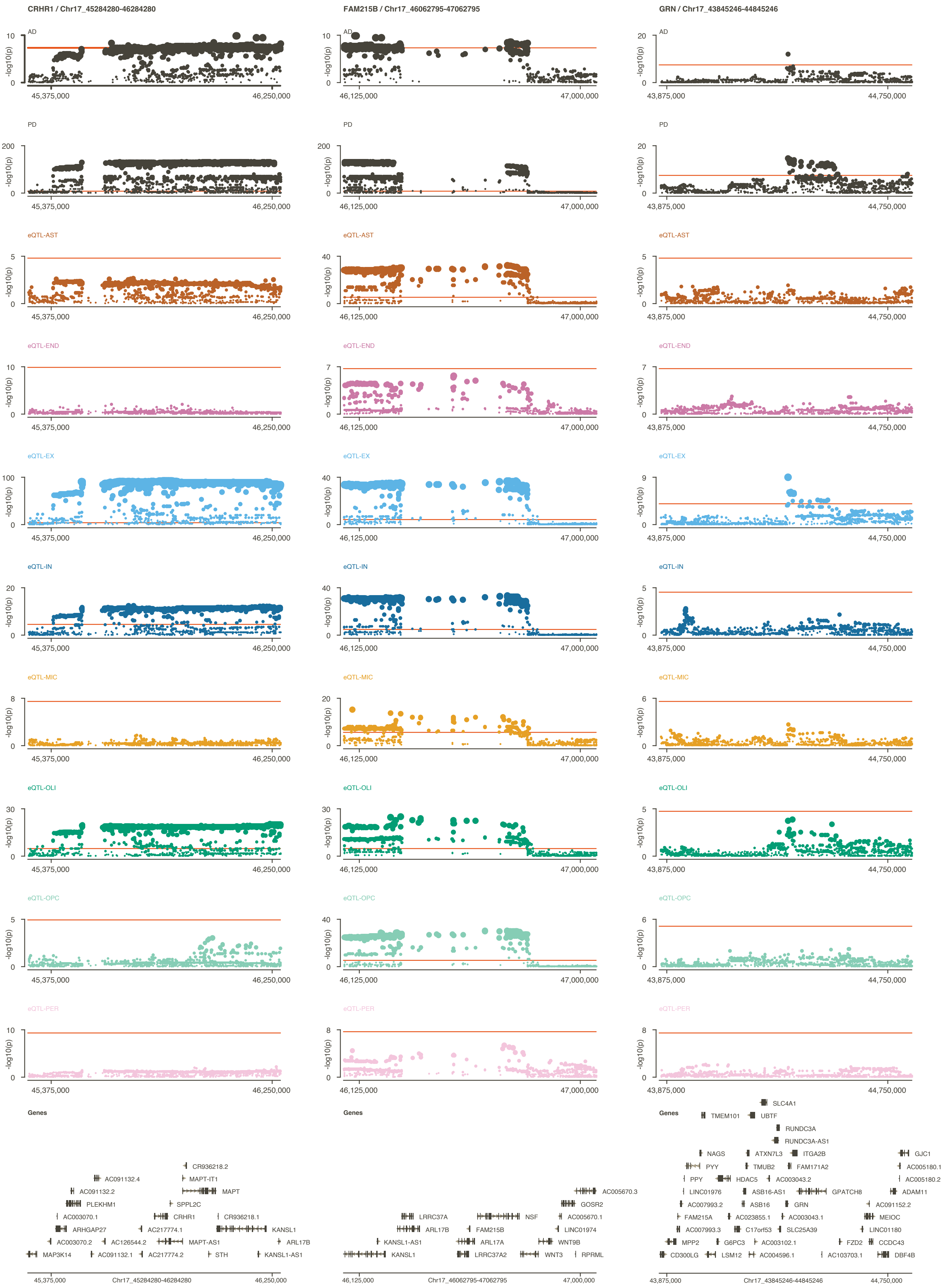

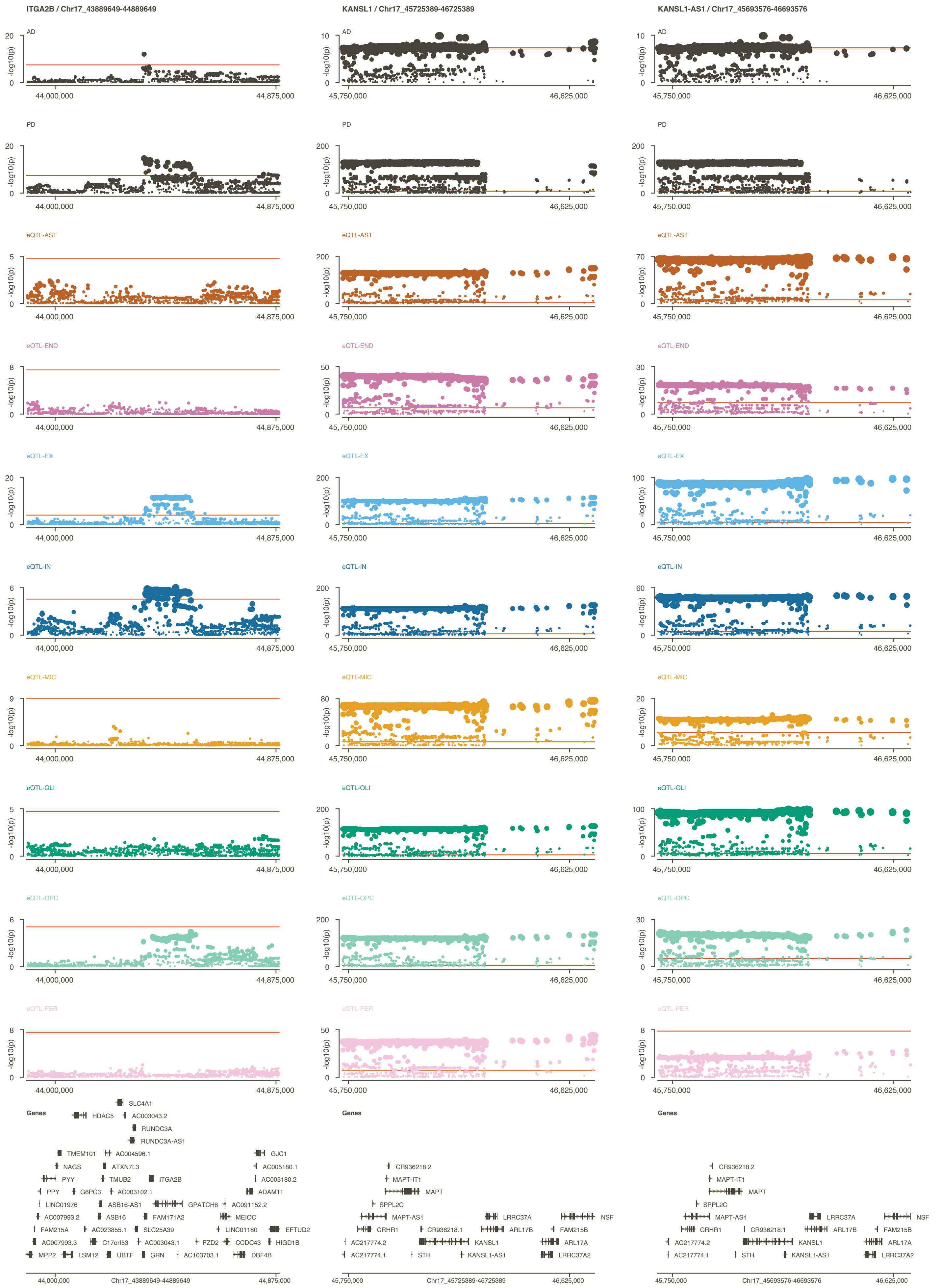

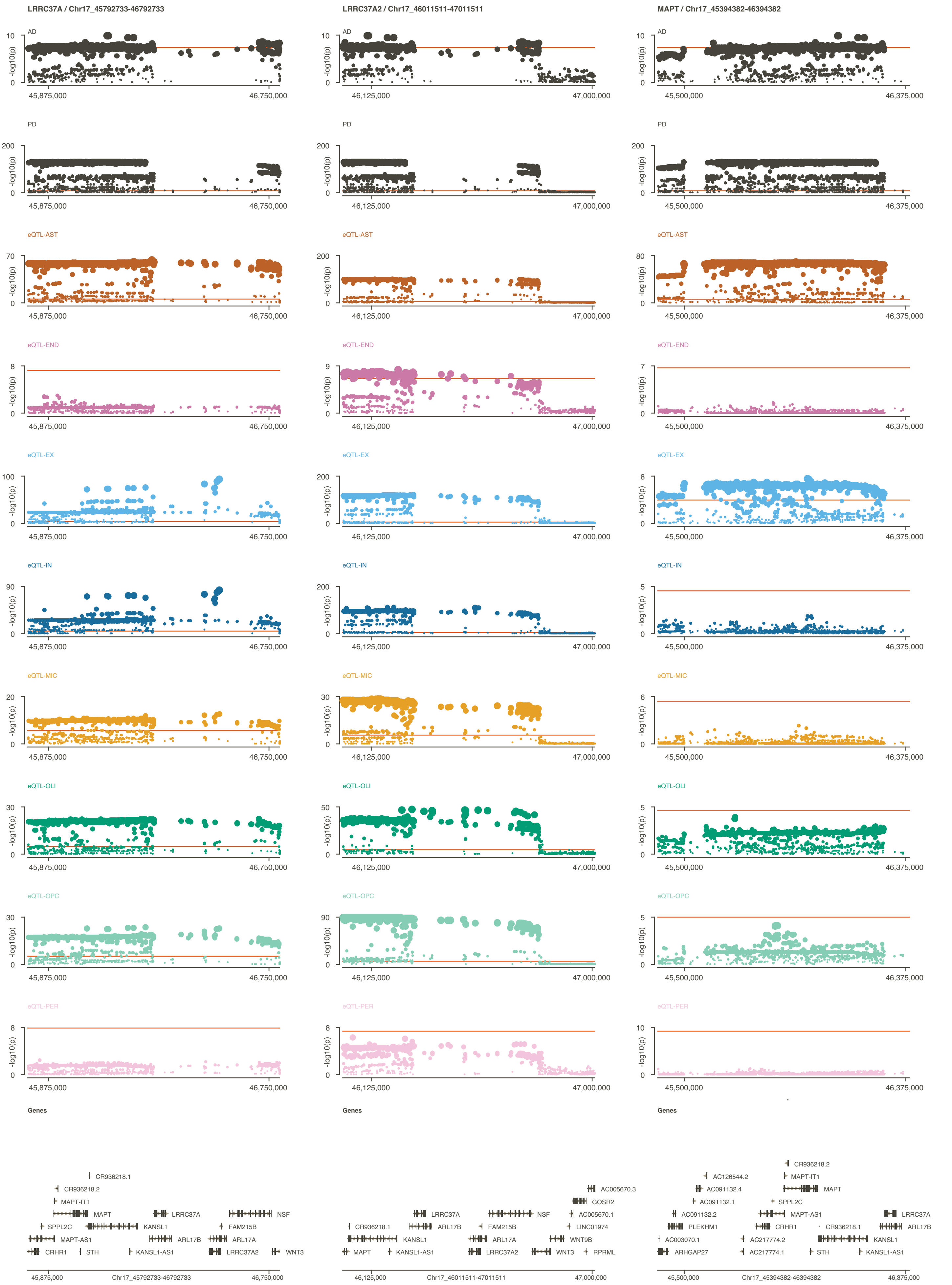

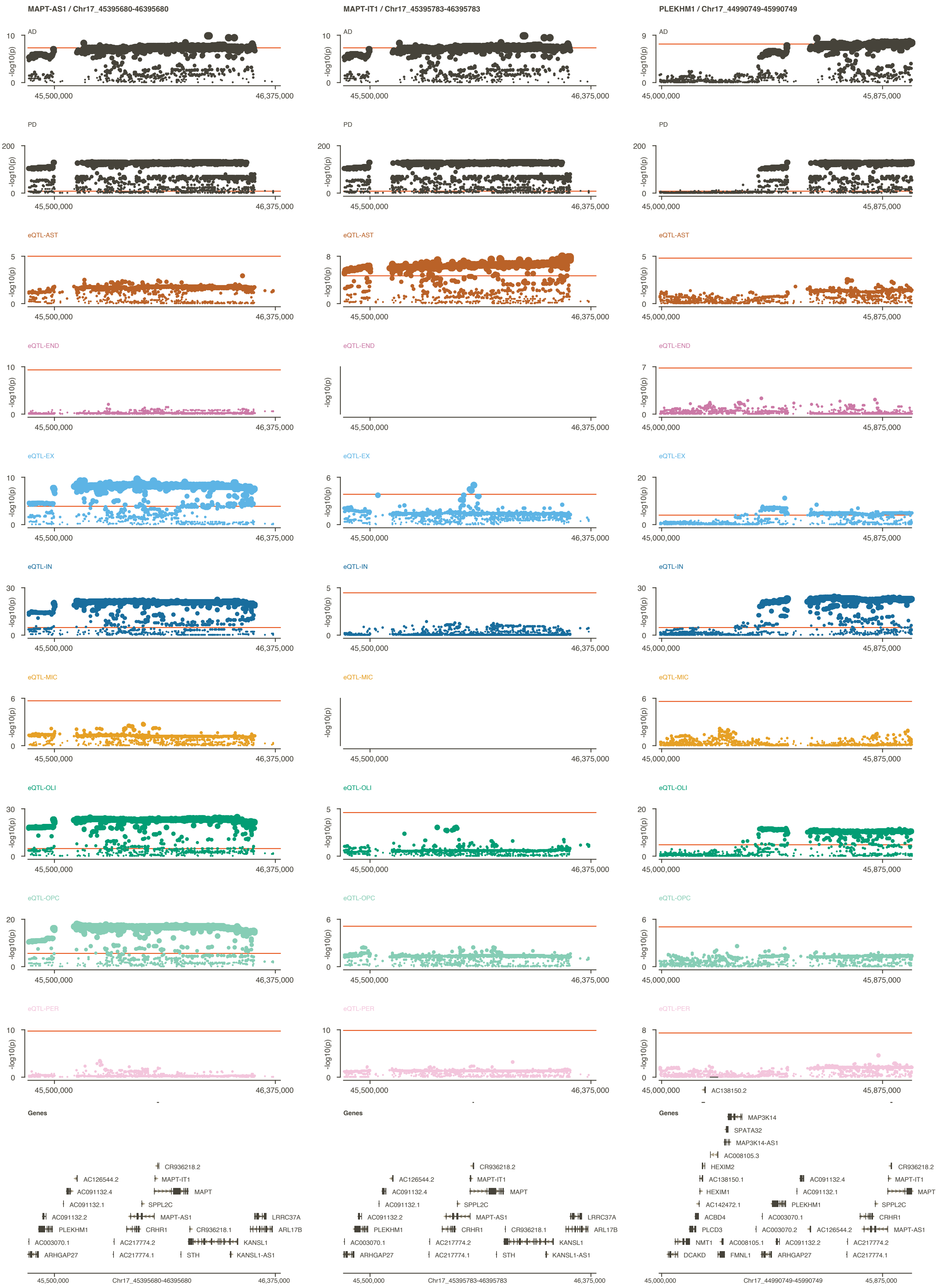

### Supplementary Figure 3

#### Supplementary Figure 3

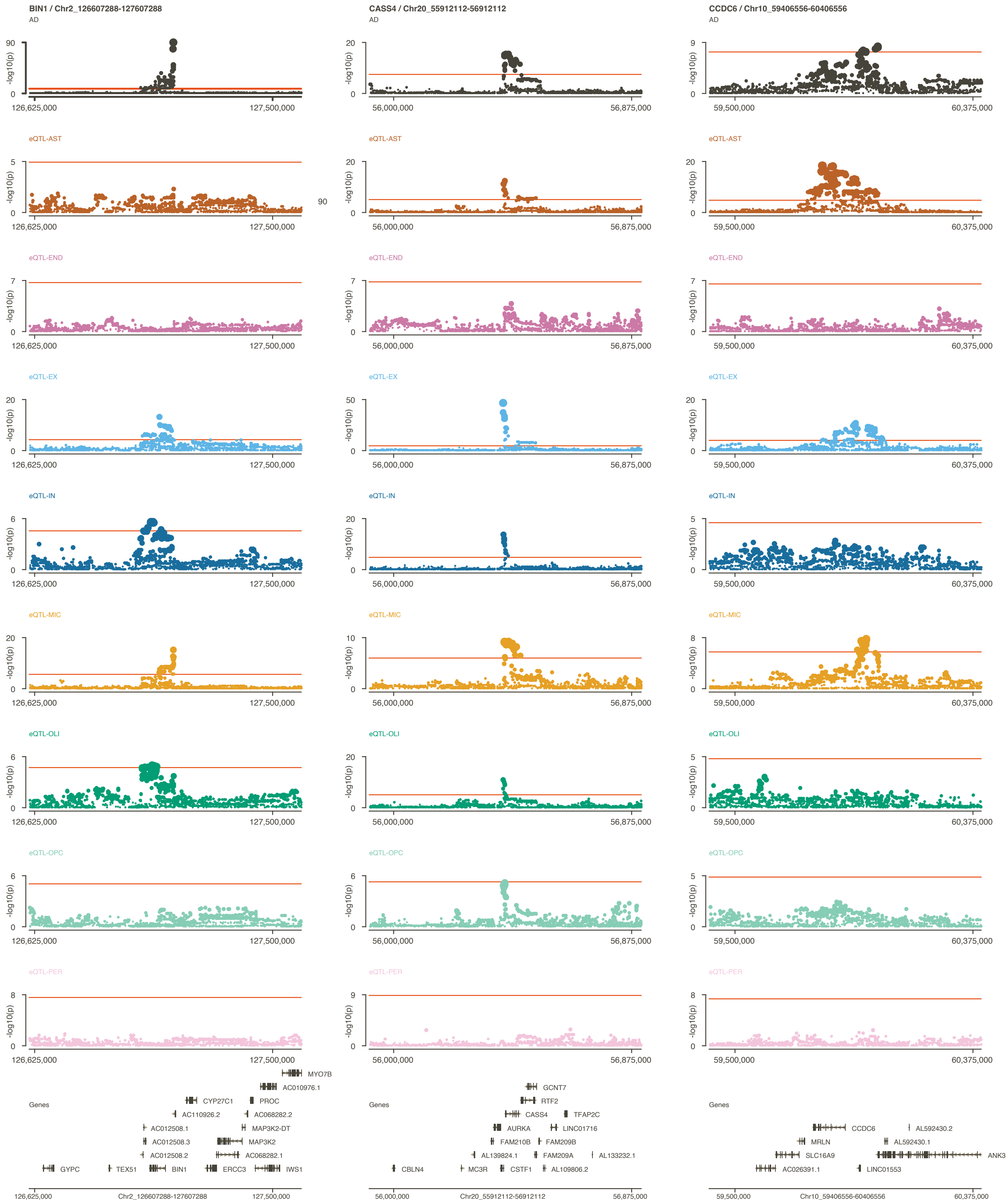

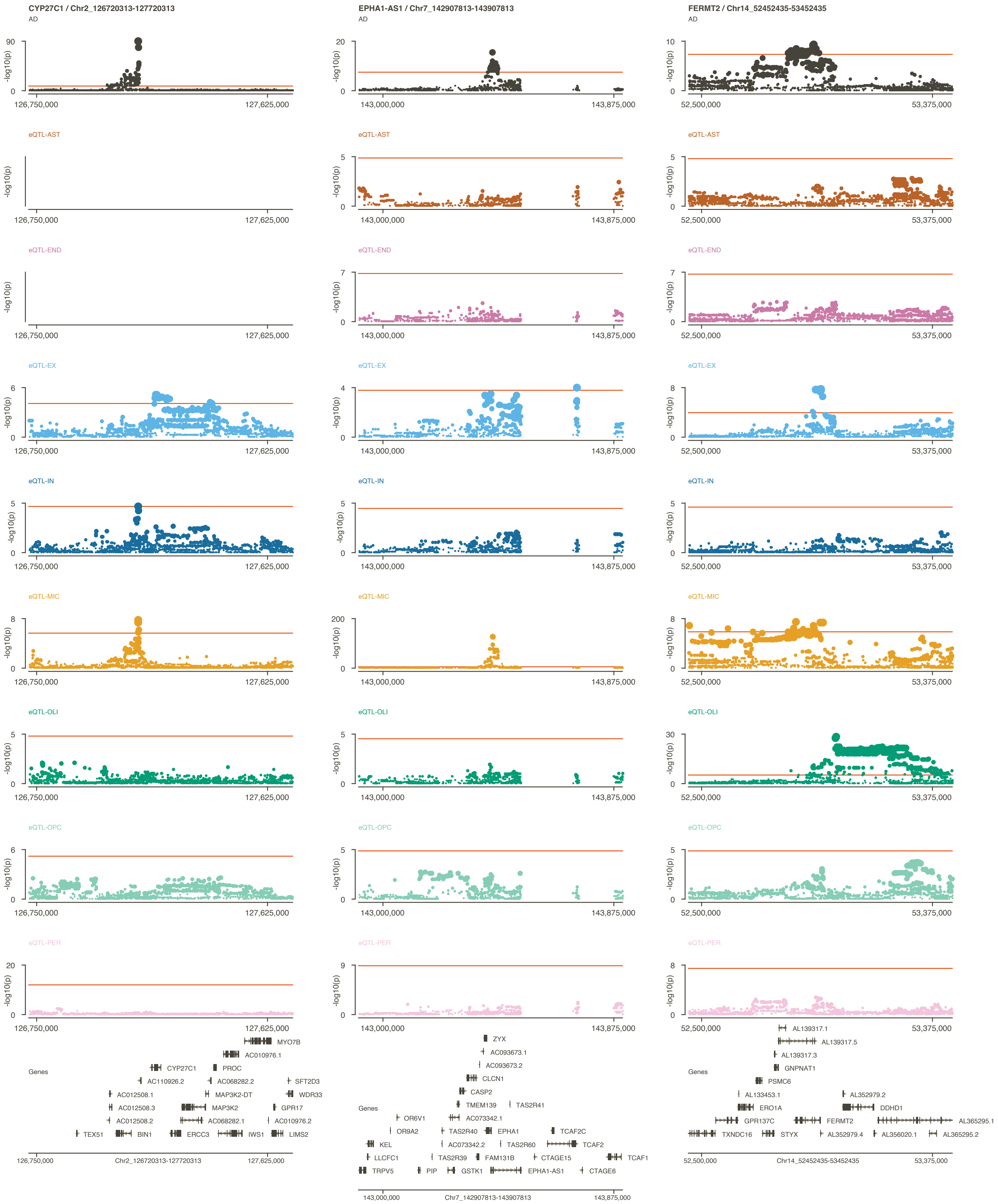

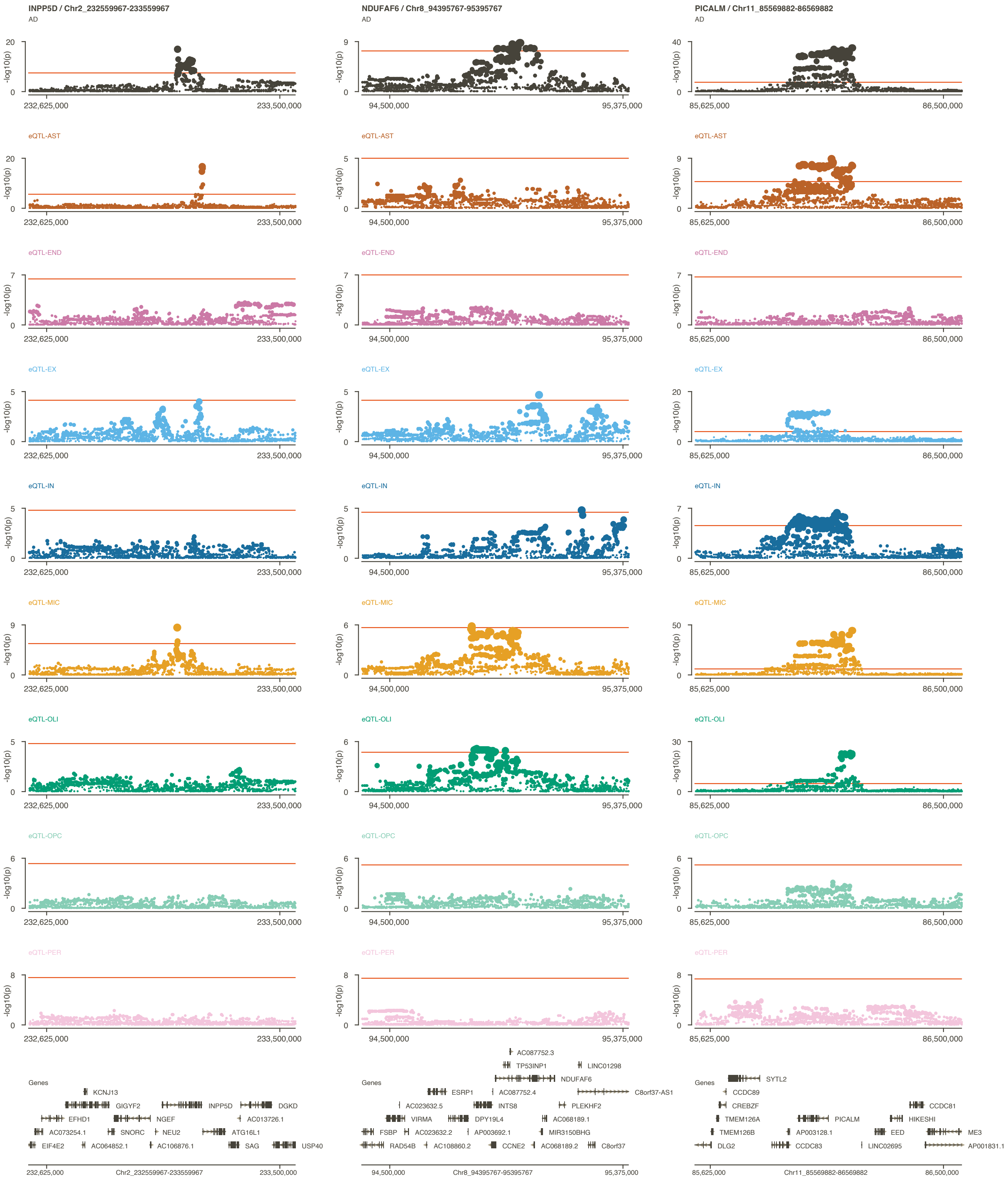

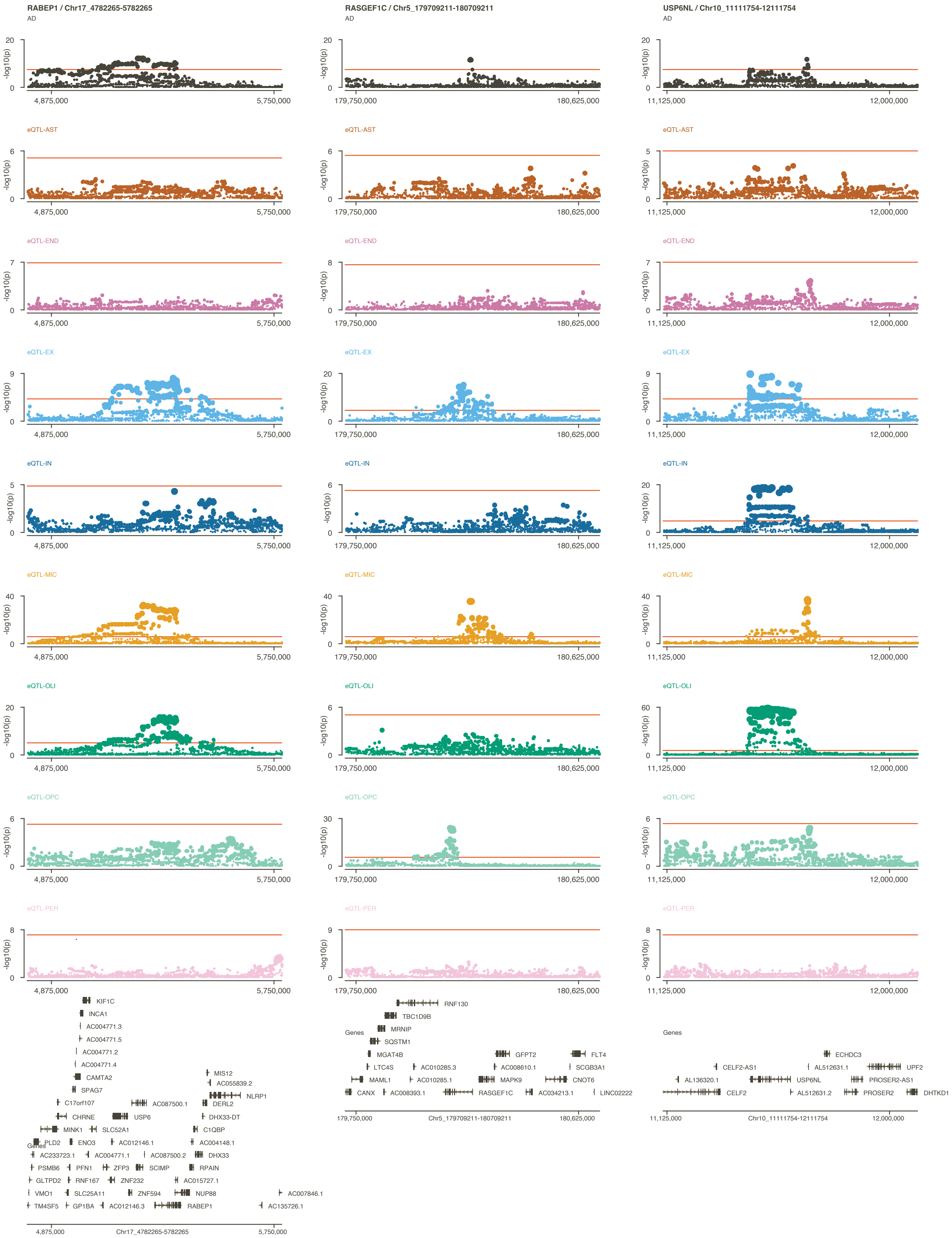

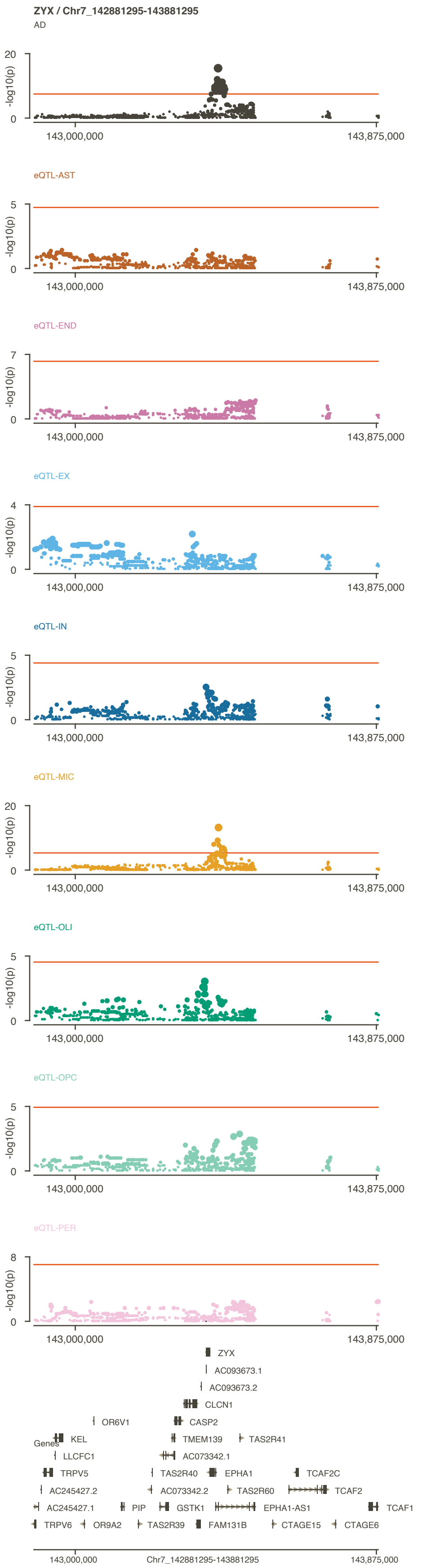
